# Mitochondrial Protein Carboxyl-Terminal Alanine-Threonine Tailing Promotes Human Glioblastoma Growth by Regulating Mitochondrial Function

**DOI:** 10.1101/2024.05.15.594447

**Authors:** Bei Zhang, Ting Cai, Esha Reddy, Yuanna Wu, Isha Mondal, Yinglu Tang, Adaeze Scholastical Gbufor, Jerry Wang, Yawei Shen, Qing Liu, Raymond Sun, Winson S Ho, Rongze Olivia Lu, Zhihao Wu

## Abstract

The rapid and sustained proliferation of cancer cells necessitates increased protein production, which, along with their disrupted metabolism, elevates the likelihood of translation errors. Ribosome-associated quality control (RQC), a recently identified mechanism, mitigates ribosome collisions resulting from frequent translation stalls. However, the precise pathophysiological role of the RQC pathway in oncogenesis remains ambiguous. Our research centered on the pathogenic implications of mitochondrial stress-induced protein carboxyl-terminal alanine and threonine tailing (msiCAT-tailing), a specific RQC response to translational arrest on the outer mitochondrial membrane, in glioblastoma (GBM). The presence of msiCAT-tailed mitochondrial proteins was observed commonly in glioblastoma stem cells (GSCs). The exogenous introduction of the mitochondrial ATP synthase F1 subunit alpha (ATP5α) protein, accompanied by artificial CAT-tail mimicking sequences, enhanced mitochondrial membrane potential (ΔΨm) and inhibited the formation of the mitochondrial permeability transition pore (MPTP). These alterations in mitochondrial characteristics provided resistance to staurosporine (STS)-induced apoptosis in GBM cells. Consequently, msiCAT-tailing can foster cell survival and migration, whereas blocking msiCAT-tailing via genetic or pharmacological intervention can impede GBM cell overgrowth.

**Impact Statement:** The Carboxyl-Terminal Alanine-Threonine-tailed protein ATP5α helps glioblastoma mitochondria maintain a high membrane potential and keep the permeability transition pore closed, thereby promoting tumor growth and increasing resistance to apoptosis.

**Highlights:**

- Glioblastoma (GBM) cells have a disturbed RQC pathway
- msiCAT-tailing on ATP5α in GBM cells increases mitochondrial membrane potential
- This msiCAT-tailing prevents MPTP opening
- ATP5α msiCAT-tailing also inhibits drug-induced apoptosis in GBM cells
- Blocking msiCAT-tailing impedes the overall growth of GBM cells

## Introduction

Proteins are vital to biological processes, and their overproduction is particularly crucial for rapidly proliferating cells, such as those found in cancer. To cope with this increased demand, cancer cells extensively reform the initiation, elongation, and termination phases of their protein synthesis (1). However, heightened protein translation elevates the chance of errors (2). Coupled with metabolic perturbations such as energy fluctuations and redox imbalances, the capacity to address disruptions during translation becomes indispensable. Ribosome-associated quality control (RQC) is a recently discovered suite of rescue mechanisms in eukaryotes that detect and resolve stalled, decelerated, or collided ribosomes during translation elongation or termination (3, 4).

RQC is a multi-step process initiated by the ZNF598/RACK1 complex, which recognizes the distinctive 40S-40S interface on collided ribosomes, triggering the ubiquitination of specific 40S subunit proteins (5, 6). Subsequently, the ASC-1 complex separates the leading ribosome (7, 8). Following this, events that transpire include: ribosomal subunit dissociation and recycling (9), modification of the nascent peptide chains by C-terminal alanine and threonine addition (CAT-tailing) (10), release of CAT-tailed products from the 60S subunits by ANKZF1/VMS1 (11), and degradation of aberrant peptides by the Ltn1/VCP/NEMF complex (4). The functional significance of CAT-tailed proteins produced during RQC remains incompletely understood. They may facilitate Ltn1-mediated ubiquitination (12) and promote the degradation of defective nascent peptides by exposing lysine residues (13, 14). Nonetheless, they are also prone to forming detergent-insoluble aggregates (15, 16). Furthermore, contingent upon the nature of the original protein and its subcellular location, CAT-tailed proteins might possess specific, albeit currently unclear, functions. Notably, CAT-tailed proteins have been implicated in the pathogenesis of several neurodegenerative diseases, indicating a significant role in their progression (17–19).

Cancerous cells exhibit increased translation irregularities, including stop codon readthrough (20), frame-shifting (21), and oxidative stress-induced ribosomal arrest (22), which suggests a potential role for the RQC pathway. While CAT-tail modification of mitochondrial proteins due to compromised RQC has been noted in HeLa cells, the mechanistic involvement of RQC factors in cancer biology remains largely unexplored (17). Notably, the expression profile of various RQC factors (e.g., ASCC3, ABCE1, ANKZF1, and VCP) is dysregulated in cancer (23–26). Interestingly, RQC factors can display opposing functions in cancer development and suppression depending on specific circumstances, with some factors like ABCE1, ASCC3, and VCP suppressing cancer cell growth upon downregulation (23, 24, 26), while others like NEMF/Clbn and ZNF598 may promote it upon inhibition (27, 28). This suggests a nuanced, context-dependent role for RQC components in cancer cells, influenced by both genetic and environmental factors. A recent study investigated the mechanism of ANKZF1 in mitochondrial proteostasis and its impact on glioblastoma multiforme (GBM) progression (29). However, this study employed a non-physiological mitochondria-targeted GFP to induce matrix proteotoxicity, leaving the role of endogenous mitochondrial proteins in this process ambiguous.

Mitochondrial stress leads to co-translational import anomalies, eliciting widespread CAT-tailing (mitochondrial stress-induced CAT-tail or msiCAT-tail) of nuclear-encoded mitochondrial proteins, including C-I30 (Complex-I 30 kDa subunit protein, NDUS3) (17, 30). The functional ramifications of these msiCAT-tailed proteins in mitochondrial biology remain poorly elucidated. Given that CAT-tailing imparts new properties to proteins, it may contribute to the distinctive features of cancer cell mitochondria, such as hyperpolarization (31, 32) and resistance to drug-induced apoptosis linked to a high mitochondrial membrane potential (Δψm) (33–35). This membrane potential across the inner membrane of mitochondria, essential for ATP production by OXPHOS, is sustained by the electron transport chain (Complexes I to IV), which pumps protons (H^+^) into the intermembrane space (36), and ATP synthase (Complex V), which leverages this gradient (37). While numerous malignant cells exhibit reduced OXPHOS despite high energy demands (38), the mechanisms by which they maintain or elevate ΔΨm remain an unresolved question (31).

In this study, we investigated msiCAT-tailing modification on the mitochondrial ATP synthase F1 subunit alpha (ATP5α). We discerned that msiCAT-tailed ATP5α is present in GBM. The mimic short-tailed ATP5α (ATP5α-AT3 in subsequent studies) can integrate into the ATP synthase, leading to an augmented ΔΨm and attenuated mitochondrial permeability transition pore (MPTP) assembly and opening. Consequently, msiCAT-tailed ATP5α enhances GBM cell resistance to programmed cell death induced by staurosporine (STS) and temozolomide (TMZ), thereby fostering cancer cell survival, proliferation, and migration. Conversely, impeding msiCAT-tailing diminishes cancer cell growth and resensitizes GBM cells to apoptosis. Our findings underscore the involvement of CAT-tailed mitochondrial proteins in tumorigenesis and emphasize the significance of the RQC pathway in oncobiology. These outcomes suggest that components and products of the RQC pathway may offer promising therapeutic targets for GBM.

## Results

### Presence of msiCAT-tailed Proteins in Glioblastoma Cells

While dysregulation of individual ribosome-associated quality control (RQC) factors is documented across various cancers (e.g., adenocarcinoma, non-small cell lung, prostate, and colon carcinomas), a comprehensive analysis of the RQC pathway in glioblastoma (GBM) has been lacking (23–26). Our analysis of transcriptomic data from a cohort of 153 GBM patients and 206 healthy controls, sourced from public datasets, revealed significantly elevated expression (logFC (fold change) > 1; adj.P.Val < 0.001) of RQC pathway genes, such as *ABCE1*, *ASCC1-3*, *RACK1*, and *VCP,* in GBM cells(39). Conversely, *ANKZF1* was significantly downregulated (logFC =-0.43, adj.P.Val = 0.0005) (Figure 1A, Table 1). The expression change in these genes implies activation of the RQC pathway and potential accumulation of CAT-tailed proteins in GBM. Mitochondrial stress-induced protein mitochondrial Complex-I 30 kDa (C-I 30, also known as NDUS3), an endogenous RQC substrate with msiCAT-tails, was previously identified in HeLa cells (17). Examination of patient-derived GBM stem cells (GSCs) and normal neural stem cells (NSCs) revealed that GSCs, unlike NSCs, exhibited several msiCAT-tailed mitochondrial proteins, including NDUS3, COX4 (Cytochrome c Oxidase subunit 4), and ATP5α (ATP synthase F1 subunit alpha). Consistent with the detection of these msiCAT-tailing signals, increased NEMF (Nuclear Export Mediator Factor) levels (10) and decreased ANKZF1 (Ankyrin Repeat and Zinc-finger Peptidyl tRNA Hydrolase 1) expression (11) were observed in patient-derived GSCs (Figure 1B), further indicative of enhanced CAT-tailing activation, mirroring bioinformatics findings in GBM samples. A murine GBM model exhibited analogous RQC pathway alterations, with increased NEMF and decreased ANKZF1 expression in transplanted SB28 gliomas compared to normal brain tissue (Figure 1 – Figure Supplement 1A, B).

**Figure 1.**
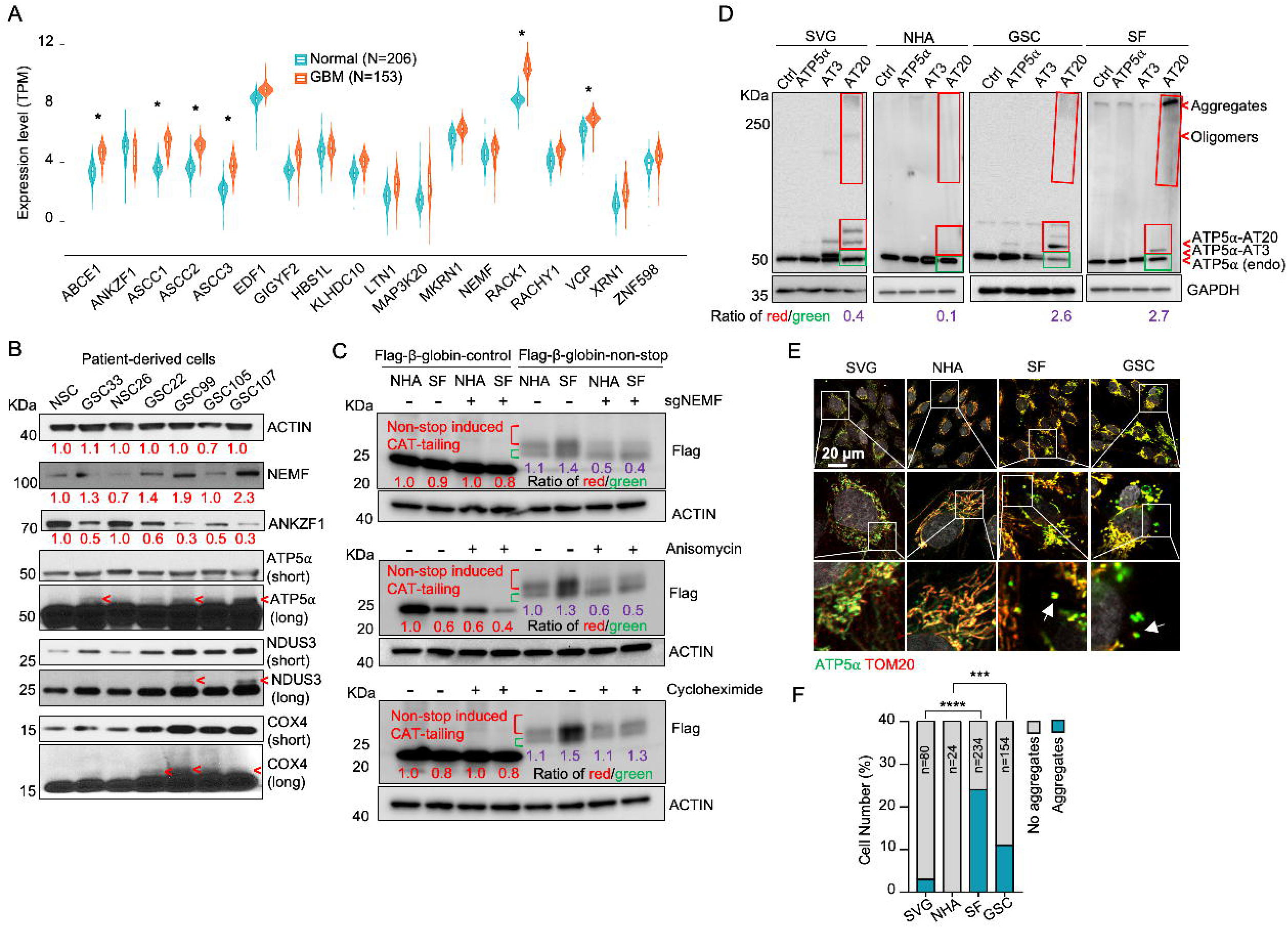
Evidence for msiCAT-tailing on mitochondrial proteins in GBM cells. (A) RQC gene expression levels in GBM tumor tissues (n=153) compared to normal brain tissues (n=206) (unpaired Student’s t-test; *, logFC (fold change)>1; adj.P.Val<0.001). (B) Western blot analysis of msiCAT-tailed mitochondrial proteins and RQC factors in patient-derived GSC and control NSC cells, using ACTIN as the loading control. Red arrowheads indicate short CAT-tailed mitochondrial proteins; “short” and “long” refer to exposure time; the red numbers represent fold changes compared to controls (NSC). (C) Western blot of 5×FLAG-tagged β-globin reporter proteins in GBM and control cells, showing more CAT-tailed proteins in GBM cells, using ACTIN as the loading control. The red numbers represent fold changes compared to controls (NHA without any treatment); the purple numbers represent the ratio of red (CAT-tailed) to green (non-CAT-tailed) sections. (D) Western blot of overexpressed ATP5α-AT3 and ATP5α-AT20 in GBM and control cells, using GAPDH as the loading control; arrowheads indicate endogenous ATP5α, ATP5α-AT3, ATP5α-AT20, and oligomers/aggregates of msiCAT-tailed ATP5α proteins. The purple numbers represent the ratio of red (exogenous) to green (endogenous) sections. (E) Immunofluorescence staining shows endogenous ATP5α protein aggregates in GBM cells, with TOM20 (red) as a mitochondrial marker. White arrows indicate ATP5α protein aggregates. (F) Quantification of E (n=3; chi-squared test; ***, *P* < 0.001; ****, *P* < 0.0001); the total number of cells counted is indicated in the columns.

**Table 1.**
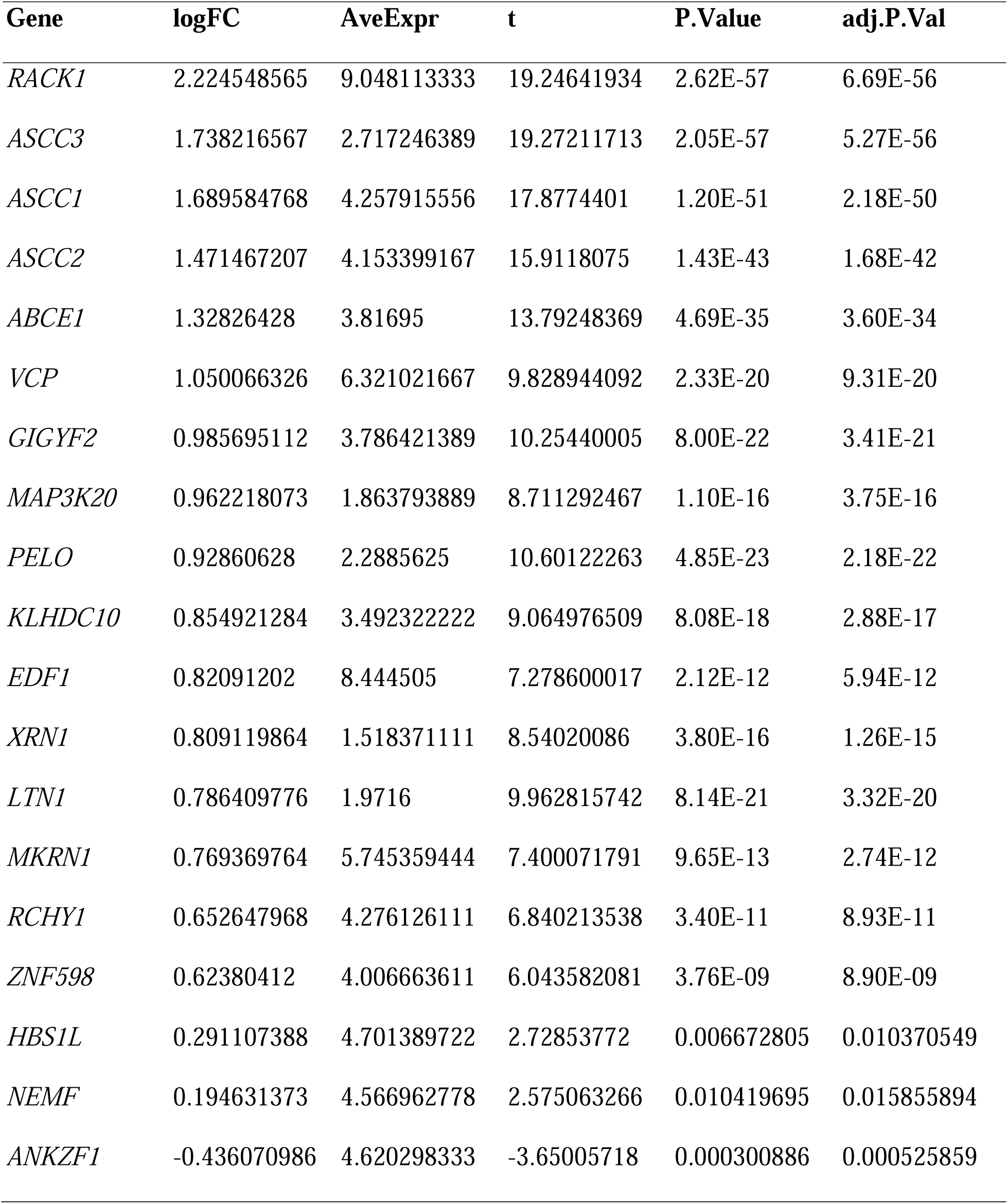
Differential expression analysis of RQC genes in GBM patients compared to healthy controls.

The subsequent experiments were conducted using two GBM cell lines, SF268 (SF in figures) (40) and GSC827 (GSC in figures) (41), and two control cell lines, SVG p12 (SVG in figures) and Normal Human Astrocytes E6/E7/hTERT (NHA in figures) (42). RQC protein expression analysis revealed decreased ANKZF1 and increased ABCE1, ASCC3, and NEMF expression in GSC827 and SF268 cells, consistent with findings in patient-derived GSCs (Figure 1 – Figure Supplement 1C). Intriguingly, induction of CAT-tailing on a Flag-tagged β-globin reporter via a non-stop protein translation system demonstrated significantly higher CAT-tailed protein (β-globin-nonstop) production in GBM cells (43). This process was inhibitable by the CAT-tailing elongation inhibitor anisomycin and NEMF knockdown (sgNEMF), but not cycloheximide treatment, as evidenced by a decreased ratio of CAT-tailed (Red) to non-CAT-tailed bands (Green) (Figure 1C).

Next, to investigate the biological implications of CAT-tailing, artificial CAT-tails were introduced to mitochondrial proteins. Due to the variability in CAT-tailing, prior research simulated this process by adding alanine-threonine (AT) repeat tails to the C-terminus of mitochondrial proteins (17). According to recent studies, the chosen tail sequence can be stabilized by its high threonine content (44, 45). ATP5α, a highly abundant mitochondrial protein with roles in cancer, was selected to study the unique functions of CAT-tailed forms (46, 47). siATP5α knockdown first confirmed the upper band signal in GSCs as authentic ATP5α, demonstrated by its disappearance concurrent with the main band’s weakening (Figure 1 – Figure Supplement 2A). Then, we confirmed that this upper band signal corresponded to changes in CAT-tailing, which could be effectively inhibited by NEMF knockdown and anisomycin treatment (Figure 1 – Figure Supplement 2B). Due to the indistinct nature of the endogenous msiCAT-tailed ATP5α signal, exogenously expressed Flag-ATP5α was utilized here.

To investigate the potential new function provided by CAT-tailed proteins, control (SVG and NHA) and GBM (SF268 and GSC827) cell lines overexpressed ATP5α with three (ATP5α-AT3) or twenty (ATP5α-AT20) AT repeats. Consistent with earlier findings, only the long-tailed ATP5α-AT20 exhibited post-translational modifications and detergent-resistant insoluble aggregates, appearing as slower migrating bands and a high molecular weight smear in protein electrophoresis (Figure 1D). Based on comparing exogenously expressed (indicated by red boxes) to endogenous proteins (indicated by green boxes), GBM cell lines (GSC827, SF268) showed increased accumulation of ATP5α-AT20 compared to control cells (SVG, NHA). This accumulation may occur due to increased stability and reduced degradation of long-tailed proteins, a malfunctioning protein quality control system, enhanced cellular tolerance to protein accumulation, or a combination of these factors. Subcellular localization analysis showed that the short AT tail (AT3) did not significantly alter ATP5α’s mitochondrial localization, similar to the tailless protein. However, a significant portion of ATP5α-AT20 was found in the cytoplasm near mitochondria, forming protein aggregates, with the highest proportion in highly malignant GSC cells (Figure 1 – Figure Supplement 2C, D). Notably, poly-glycine-serine tails (short, GS3, and long, GS20) did not induce insoluble protein aggregation or intracellular punctate distribution (Figure 1 – Figure Supplement 2E-G), highlighting the importance of specific amino acid composition.

Importantly, in GBM cells, both exogenous tailed proteins and the endogenous ATP5α formed clusters attached to the outer mitochondrial membrane (Figure 1E, F). Similar aggregate formation in GBM cells was also observed with other mitochondrial proteins, such as NDUS3 (Figure 1 – Figure Supplement 3A, B). Furthermore, we examined the mouse GBM models. Akin to *in vitro* culture, ATP5α in transplanted SB28 glioma formed more punctate signals and did not always colocalize with the mitochondrial marker TOM20 (Figure 1 – Figure Supplement 3C-E). These findings collectively indicate a disruption of the RQC pathway, leading to the presence of msiCAT-tailed proteins in GBM cells.

### msiCAT-tailed ATP5***α*** Elevates Mitochondrial Membrane Potential (***ΔΨ***m)

Some cancer cells exhibit altered mitochondrial physiology, maintaining or increasing mitochondrial membrane potential (ΔΨm) despite reduced respiration. This was observed in patient-derived GSC cells, which demonstrated higher ΔΨm but lower ATP production than control NSC cells (Figure 2A, B). Similarly, GBM cell lines, GSC827 and SF268, displayed comparable or higher ΔΨm and lower ATP levels relative to the control NHA cell line (42) (Figure 2 – Figure Supplement 1A-C). Genetic inhibition of msiCAT-tailing, via NEMF knockdown (sgNEMF) or ANZKF1 overexpression (oeANZKF1) (Figure 2 – Figure Supplement 1D), as well as pharmacological inhibition by anisomycin treatment, effectively reduced ΔΨm in GBM cells but not in NHA cells (Figure 2C, D).

**Figure 2.**
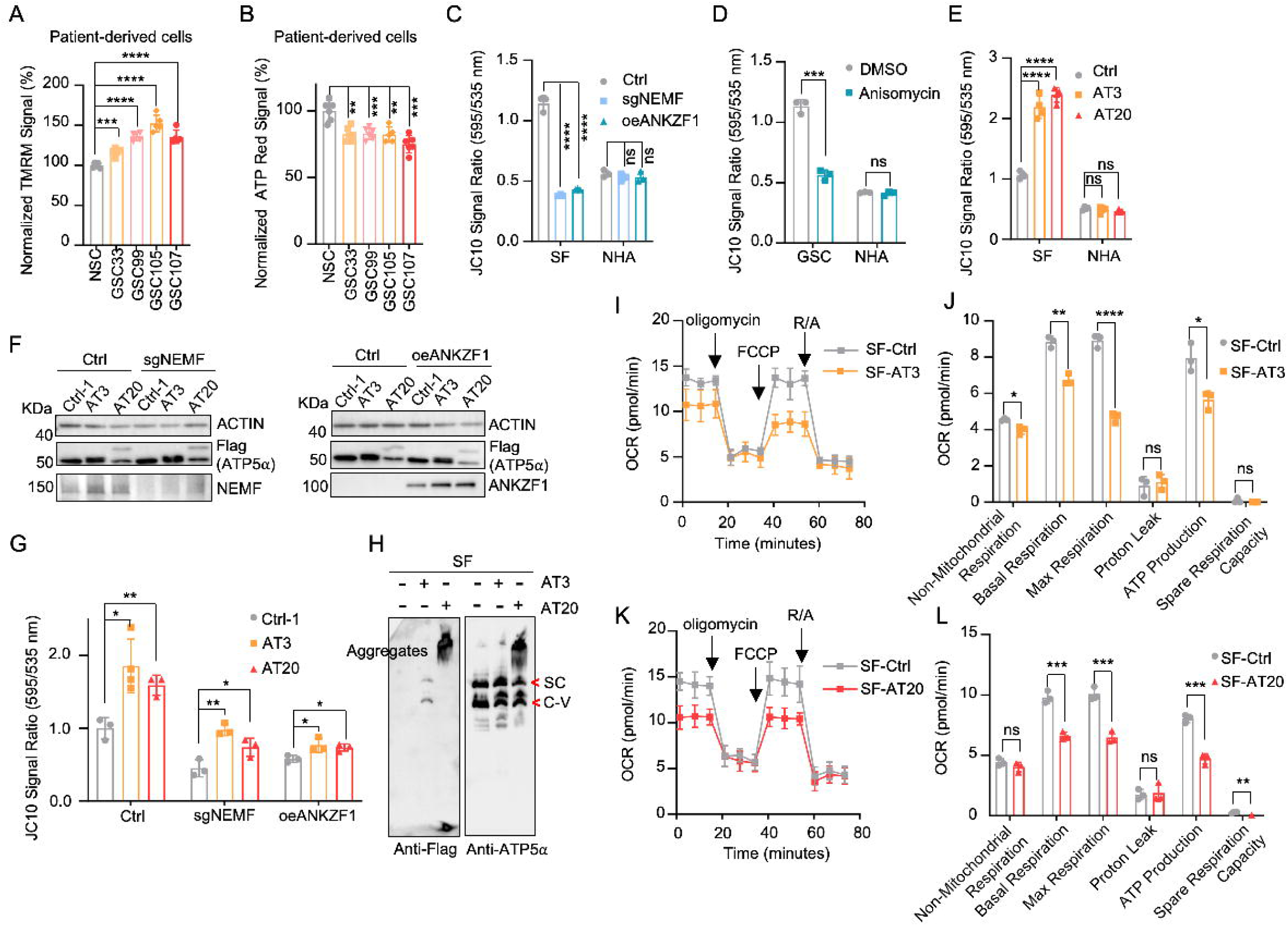
Impact of msiCAT-tailed ATP5IZ proteins on mitochondrial functions in GBM cells. (A) TMRM staining shows a high mitochondrial membrane potential in patient-derived GSC cells (n=3; unpaired Student’s t-test; ***, *P* < 0.001; ****, *P* < 0.0001). (B) ATP measurement shows a low mitochondrial ATP production in patient-derived GSC cells (n=3; unpaired Student’s t-test; \**\**, *P* < 0.01; ***, *P* < 0.001). (C, D) JC-10 staining reveals a reduced mitochondrial membrane potential in GBM cells, but not in NHA control cells, upon both genetic (C) and pharmacological (D) inhibition of the msiCAT-tailing pathway (n=3; unpaired Student’s t-test; ***, *P* < 0.001; ****, *P* < 0.0001; ns, not significant). (E) JC-10 staining reveals an increased mitochondrial membrane potential in GBM cells, but not in control cells, upon overexpression of ATP5⍰-AT3 and ATP5⍰-AT20 (n=3; unpaired Student’s t-test; ****, *P* < 0.0001; ns, not significant). (F) Western blot of FLAG-tagged ATP5⍰, NEMF, and ANKZF1 in GBM cells and control cells, using ACTIN as the loading control. (G) JC-10 staining reveals an increased mitochondrial membrane potential in GBM cells, but not in NHA control cells, upon overexpression of ATP5⍰-AT3 and ATP5⍰-AT20 with concurrent genetic inhibition of the endogenous msiCAT-tailing pathway (n=3; unpaired Student’s t-test; *, *P* < 0.05; **, *P* < 0.01). (H) BN-PAGE western blot of ATP5⍰ and Flag shows that ATP5⍰-AT3 is incorporated into the mitochondrial Complex-V (ATP synthase), while ATP5⍰-AT20 forms high molecular weight protein aggregates in GBM cells. SC: respiratory supercomplex; C-V: Complex-V/ATP synthase. (I, K) Oxygen consumption rate (OCR) data indicate a reduction in mitochondrial oxygen consumption in SF268 cells expressing ATP5⍰-AT3 and ATP5⍰-AT20. Oligomycin (1.5 µM), FCCP (1.0 µM), and rotenone/antimycin A (R/A, 0.5 µM) were sequentially added. (J, L) Statistics of mitochondrial respiration parameters in (I, K), including non-mitochondrial respiration, basal respiration, maximum respiration, spare respiration, proton leaks, and ATP production (n=3; unpaired Student’s t-test; *, *P* < 0.05; \**\**, *P* < 0.01; ***, *P* < 0.001; ****, *P* < 0.0001; ns, not significant).

Our next investigation of msiCAT tail proteins revealed their impact on mitochondrial function. Expression of Flag-tagged ATP517-AT3 and ATP517-AT20 in GBM and control cell lines elevated ΔΨm specifically in GBM cells (Figure 2E). Overexpression of ATP517-GS3 and ATP517-GS20 did not exhibit this effect (Figure 2 – Figure Supplement 1E). To our surprise, even with suppressed endogenous CAT-tailing through sgNEMF and oeANZKF1 in GSC cells, the introduced AT3 and AT20 proteins could still effectively elevate ΔΨm (Figure 2F, G). This finding suggests that CAT-tailing of ATP517 may be a significant contributor to the observed mitochondrial phenotype (Figure 2G). Blue Native Polyacrylamide Gel Electrophoresis (BN-PAGE) illustrated distinct effects based on CAT-tail length. ATP517-AT3 integrated into the mitochondrial respiratory chain complex, whereas ATP517-AT20 formed high molecular weight complexes or remained as monomers (Figure 2H). In mitochondrial physiological activity assays using the Agilent Cell Mitochondrial Stress Test, the oxygen consumption rate (OCR) was directly measured to assess mitochondrial respiration. Our findings indicate that expressing both ATP517-AT3 and ATP517-AT20 negatively impacted mitochondrial oxidative phosphorylation. This impairment leads to a reduction in ATP synthesis, basal respiration, and maximal respiration rates (Figure 2I-L). These data suggest that both short and long tails on ATP517 proteins influence mitochondrial function, although potentially through different mechanisms. Short CAT-tails may directly act on ATP synthase function and thus affect the respiratory chain complex, while long CAT-tails form protein aggregates, causing mitochondrial proteostasis stress and thus indirectly affecting mitochondrial respiration (17, 29). This differential impact of CAT-tail length suggests a nuanced regulation of mitochondrial function mediated by ATP517 modifications.

### msiCAT-tailing Influences Mitochondrial Permeability Transition Pore (MPTP) Dynamics

Beyond its traditionally recognized role in ATP production, the F_1_F_0_ ATP synthase has garnered increasing attention as a potential structural component of the mitochondrial permeability transition pore (MPTP) complex (48–50). Given the possibility that CAT-tailed proteins like ATP517 might modulate MPTP function, this investigation sought to elucidate the mechanism by which msiCAT-tailing modulates MPTP dynamics (open-close state). Comparative analyses conducted in GBM and control cells revealed that MPTP in GSC827 cells predominantly exists in a closed conformation, indicated by strong Calcein signals. Notably, the treatment of anisomycin, a pharmacological CAT-tailing inhibitor, effectively induced MPTP opening in GSC827 cells, as indicated by decreased Calcein signals (Figure 3A, B). This effect was concomitant with the diminished aggregation of endogenous ATP5α (Figure 3E, F). Furthermore, corroborative evidence was obtained through genetic manipulation. Specifically, genetic inhibition of CAT-tailing via NEMF knockdown (sgNEMF) resulted in a similar decrease in Calcein signaling and a reduction in ATP5α accumulation (Figure 3C, D, G, H), aligning with the results obtained using anisomycin. In contrast, treatment with cycloheximide, a general translation inhibitor, did not significantly alter Calcein or ATP517 aggregation signals (Figure 3 – Figure Supplement 1A-D), suggesting that non-specific translation inhibition does not impact the mitochondrial MPTP state. The crucial role of CAT-tail modifications on ATP5α in modulating MPTP status was further substantiated by the observation that overexpression of artificially synthesized AT repeat tails (AT3 and AT20) restored Calcein signals despite the inhibition of endogenous CAT-tailing (Figure 3 – Figure Supplement 1E).

**Figure 3.**
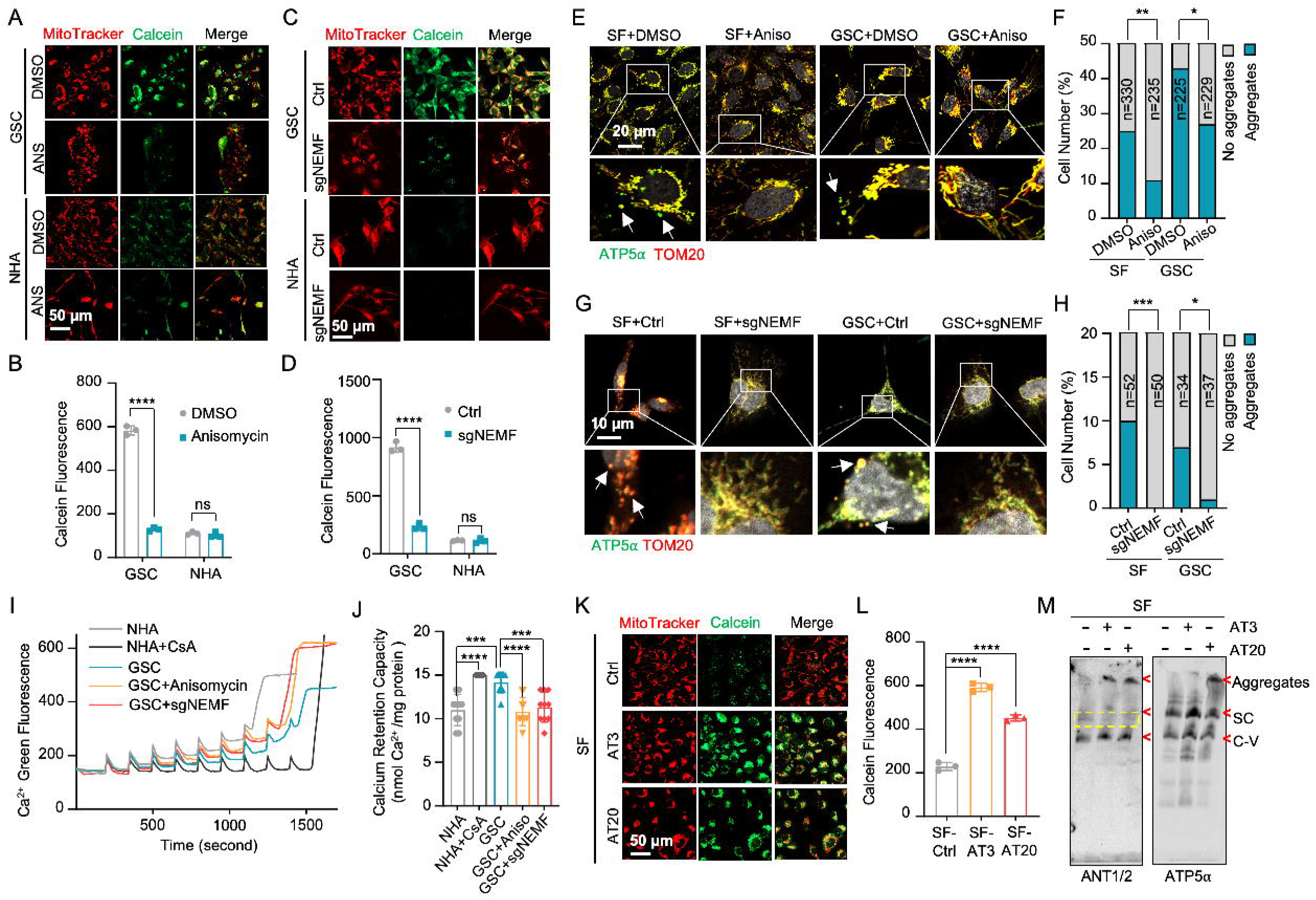
msiCAT-tailing product regulates MPTP status in GBM cells. (A, C) MPTP activity assay shows reduced mitochondrial permeability transition pore opening in GSC cells compared to NHA (control) cells. Pharmacological (A, anisomycin 200 nM) and genetic (sgNEMF) inhibition of CAT-tailing reverse it. (B, D) Quantification of (A, C) (n=3; unpaired Student’s t-test; ****, *P* < 0.0001; ns, not significant). (E, G) Immunofluorescence staining shows that anisomycin treatment (E) and sgNEMF (G) inhibit endogenous ATP5⍰ protein aggregation in GBM cells, using TOM20 (red) as a mitochondrial marker. (F, H) Quantification of (E, G) (n=3; chi-squared test; *, *P* < 0.05; **, *P* < 0.01; ***, *P* < 0.001); the total number of cells counted is indicated in the columns. (I) The calcium retention capacity (CRC) assay of isolated mitochondria, measured using the Calcium Green-5N dye, reveals a significantly higher CRC in GBM cells compared to control NHA cells. CsA (Cyclosporin A, MPTP inhibitor) serves as a positive control. (J) Statistic of (I) shows attenuated CRC in mitochondria pretreated with anisomycin or with sgNEMF (n=10; unpaired Student’s t-test; ***, *P* < 0.001; ****, *P* < 0.0001). (K) MPTP activity assay shows that ectopic expression of ATP5⍰-AT3 and ATP5⍰-AT20 inhibits MPTP opening in GBM cells. (L) Quantification of (K) (n=3; unpaired Student’s t-test; ****, *P* < 0.0001). (M) BN-PAGE western blot shows that ATP5⍰-AT3 and ATP5⍰-AT20 expression alters ANT1/2 protein patterns in GBM cells, resulting in a missing band (circled in yellow dashed line), and formation of high molecular weight aggregates. SC: respiratory supercomplex; C-V: Complex V/ATP synthase.

The MPTP is recognized to participate in the transient efflux of protons, calcium ions (Ca²□), and other signaling molecules from the mitochondrial matrix during brief opening episodes (51). To quantitatively evaluate the MPTP open/closed state, the mitochondrial Ca² Retention Capacity (CRC) assay was employed, which measures the amount of Ca² required to elicit MPTP opening. Our results revealed that GSC827 cells exhibited a greater CRC value than NHA cells. Pre-treatment with anisomycin or knockdown of NEMF (sgNEMF) significantly decreased the CRC in GBM cells, indicating MPTP opening upon the loss of CAT-tailed proteins (Figure 3I, J). Consistent with Calcein staining results (Figure 3 – Figure Supplement 1A, B), cycloheximide treatment did not substantially alter CRC measurements (Figure 3 – Figure Supplement 2A, B). Conversely, enhancing CAT-tailing (e.g., via oeNEMF and siANKZF1) led to an increase in CRC (Figure 3 – Figure Supplement 2A, B), although this effect was less pronounced in GSCs, potentially due to their inherently active CAT-tailing and closed MPTP.

To further investigate the impact of specific AT repeat tails on MPTP opening, artificial AT repeat tails on ATP517 were introduced into GBM cells. It was found that the short AT tail (AT3) inhibited MPTP opening, while the long AT tail (AT20) displayed a weaker effect (Figure 3K, L), potentially due to their different integration into ATP synthase (Figure 2G). Complex co-immunoprecipitation assay did not detect direct interactions between ATP517 with AT3 or AT20 tails and MPTP components Cyclophilin D (Cyp-D) and adenine nucleotide translocator 2 (ANT2) (Figure 3 – Figure Supplement 2C). However, Cyp-D expression was reduced upon ectopic expression of ATP517-AT3 and ATP517-AT20, suggesting decreased MPTP formation (Figure 3 – Figure Supplement 2D). Intriguingly, BN-PAGE analysis revealed that both ATP517-AT3 and ATP517-AT20 altered ANT1/2-containing complexes, with expected bands disappearing (indicated by *) and aggregates forming (at the top), supporting the notion that ATP synthase is integrated into the MPTP supercomplex due to the spatial proximity of the ANT1/2 complex and ATP synthase (Figure 3M). In conclusion, msiCAT-tailed ATP517 proteins, particularly those with short AT3 tails, are integrated into ATP synthase and have a substantial influence on modulating MPTP status.

### msiCAT-Tailing Boosts GBM Cell Migration and Resistance to Apoptosis

The elevated mitochondrial membrane potential (ΔΨm) and constricted MPTP resulting from msiCAT-tailed ATP517 and other mitochondrial proteins may enhance cellular stress resilience. We first investigated how the msiCAT-tailing mechanism affects GBM cells at the cellular level.

MTT assays (52) revealed that overexpressing short (AT3) and long (AT20) AT repeat tails, fused to ATP517, significantly improved GBM cell viability, but not that of NHA cells (Figure 4A, B). However, short (GS3) and long (GS20) GS repeat tails did not affect GBM cell viability (Figure 4 – Figure Supplement 1A). In addition, *in vitro* transwell migration assays (53) and wound healing assays (54) showed that GBM cells overexpressing AT repeat-tailed ATP5α exhibited increased cell invasion and accelerated wound healing, indicating enhanced cell migration (Figure 4C, D, Figure 4 – Figure Supplement 1B, C). Notably, neither ATP5α alone nor GS repeat-tailed proteins showed comparable changes (Figure 4 – Figure Supplement 1D, E). Furthermore, overexpressing AT3- and AT20-tailed proteins effectively conferred phenotypes associated with increased GBM cell activity, such as enhanced survival and migration, even with inhibited endogenous CAT-tailing machinery activity (e.g., sgNEMF and oeANKZF1) (Figure 4E-G). It is worth noting that ANKZF1 knockdown in U87 and U251 cell lines can cause aberrant mitoGFP accumulation, possibly reducing cellular adaptability (29), suggesting varying mitochondrial adaptability to proteostasis stress across cell lines. Supporting this, initial experiments showed that mild expression of ATP5α-AT3 and ATP5α-AT20 did not induce strong mitochondrial proteotoxic responses, as evidenced by the lack of significant upregulation in *LONP1*, *mtHSP70*, and *HSP60* mRNA levels (Figure 4 – Figure Supplement 1F).

**Figure 4.**
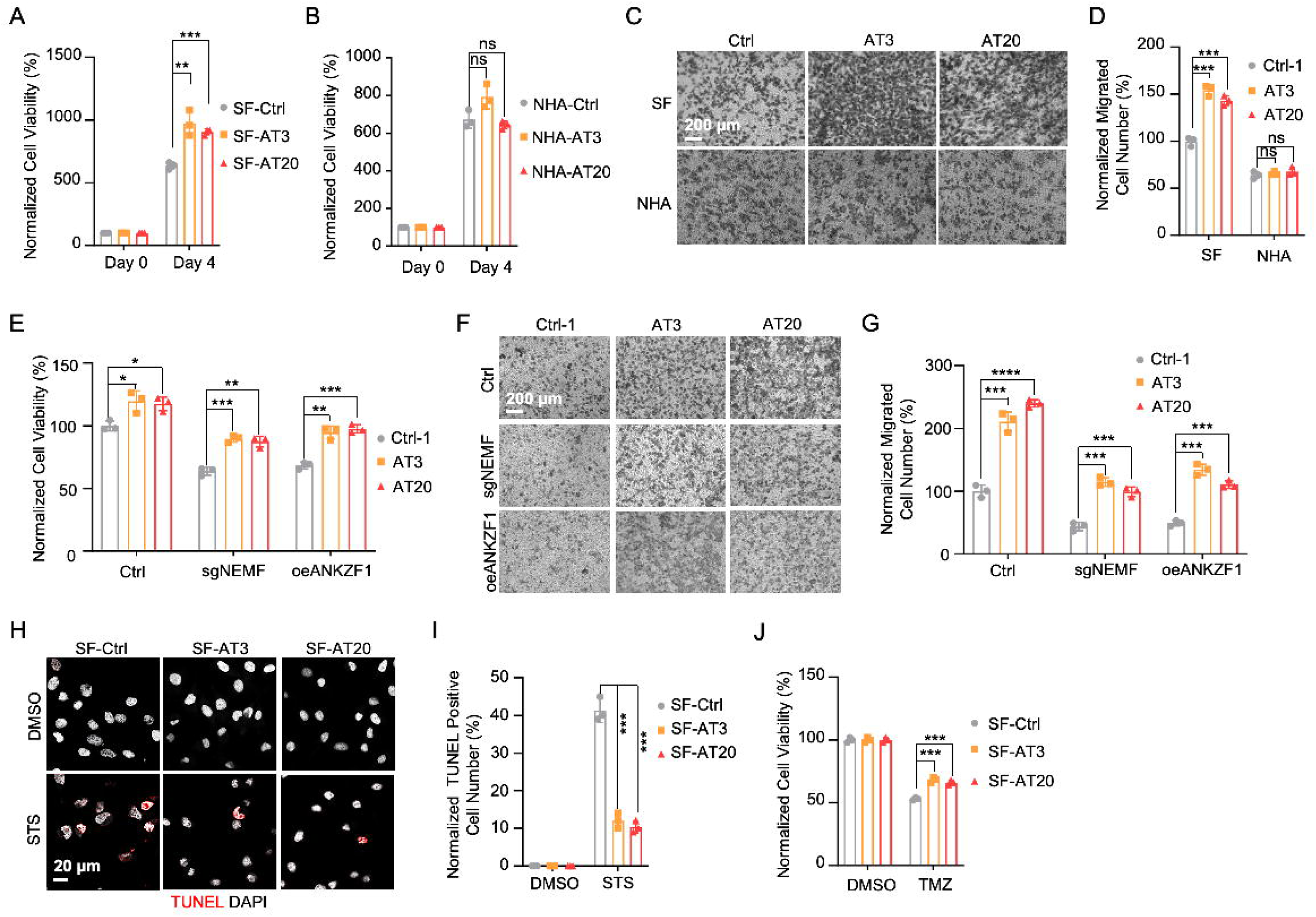
msiCAT-tailed ATP5IZ protein promotes GBM progression. (A) MTT assay indicates increased proliferation caused by ATP5⍰-AT3 and ATP5⍰-AT20 expression in GBM cells (n=3; unpaired Student’s t-test; **, *P* < 0.01; ***, *P* < 0.001). (B) MTT assay indicates no change in proliferation caused by ATP5⍰-AT3 and ATP5⍰-AT20 expression in NHA cells (n=3; unpaired Student’s t-test; ns, not significant). (C) Transwell assay reveals enhanced migration induced by ATP5⍰-AT3 and ATP5⍰-AT20 expression in GBM (SF) cells but not in control (NHA) cells. (D) Quantification of (C) shows the number of migrated cells (n=3; unpaired Student’s t-test; ***, *P* < 0.001; ns, not significant). (E) MTT assay indicates an increased proliferation in GBM cells, upon overexpression of ATP5⍰-AT3 and ATP5-AT20 with concurrent genetic inhibition of the endogenous msiCAT-tailing pathway (n=3; unpaired Student’s t-test; *, *P* < 0.05; **, *P* < 0.01). (F) Transwell assay reveals enhanced migration upon overexpression of ATP5⍰-AT3 and ATP5⍰-AT20 with concurrent genetic inhibition of the endogenous msiCAT-tailing pathway. (G) Quantification of (F) shows the number of migrated cells (n=3; unpaired Student’s t-test; ***, *P* < 0.001; ****, *P* < 0.0001). (H) TUNEL staining shows that staurosporine (STS, 1 µM)-induced apoptosis is attenuated by ATP5⍰-AT3 and ATP5⍰-AT20 expression in GBM cells, using TUNEL-Cy3 as an apoptotic cell indicator and DAPI as a nucleus indicator. (I) Quantification of (H) shows the percentage of TUNEL-positive cells in the population (n=3; unpaired student’s t-test; ***, *P* < 0.001), using DMSO as the vehicle control. (J) MTT assay indicates an enhanced resistance to temozolomide (TMZ, 150 µM) induced by ATP5⍰-AT3 and ATP5⍰-AT20 expression. The TMZ-treated/SF-Ctrl group is used as the control (n=3; unpaired Student’s t-test; ***, *P* < 0.001).

GBM cells exhibit increased resistance to staurosporine (STS)-induced apoptosis, supported by fewer TUNEL-positive cells (Figure 4 – Figure Supplement 2A, B) and markedly diminished PARP-1 (Poly ADP-ribose polymerase) cleavage (Figure 4 – Figure Supplement 2C), a marker of AIF-mediated apoptosis (55). To investigate the role of CAT-tailed ATP5α proteins in this resistance, we overexpressed proteins with mimetic tails in GBM cells. Overexpression of both short tail (ATP5α-AT3) and long tail (ATP5α-AT20) significantly enhanced resistance to STS-induced apoptosis, as shown by TUNEL staining (Figure 4H, I) and flow cytometry (Figure 4 – Figure Supplement 2D, E), indicating a strong link between protein CAT-tailing and tumorigenesis. In contrast, control short (GS3) and long (GS20) GS tails failed to confer such resistance (Figure 4 – Figure Supplement 2F, G). Consistent with these findings, overexpression of artificial CAT-tailed ATP5α proteins also increased the resistance of GBM cells to temozolomide (TMZ)-induced apoptosis (Figure 4J). Taken together, these results suggest that RQC-induced CAT-tailing on ATP5α protein plays a role in GBM resistance to drug-induced apoptosis.

### RQC Pathway Inhibition hinders GBM Cell Progression

Prior research indicates the RQC pathway’s mediated msiCAT-tailing plays an important role in GBM progression, suggesting it as a potential therapeutic target. To explore this, patient-derived Glioblastoma Stem Cell (GSC) lines were treated with anisomycin, an inhibitor of CAT-tailing. GSC lines displayed higher sensitivity to anisomycin than normal neural stem cells (NSCs) (Figure 5A). Similarly, genetic inhibition of the RQC pathway via NEMF knockdown (sgNEMF) or ANKZF1 overexpression (oeANZKF1) in the SF268 GBM cell line also suppressed GBM growth (Figure 5B). Notably, control NHA cell proliferation was also inhibited by these genetic changes, indicating the broad significance of NEMF and ANKZF1 in cell proliferation (Figure 5C). The RQC pathway appears to have a more pronounced effect on GBM cell migration. In *in vitro* transwell assays, sgNEMF or oeANZKF1 notably decreased GBM cell migration without affecting NHA cells (Figure 5D, E). Consistently, anisomycin treatment impaired GSC cell migration, but not NHA cell migration (Figure 5F, G).

**Figure 5.**
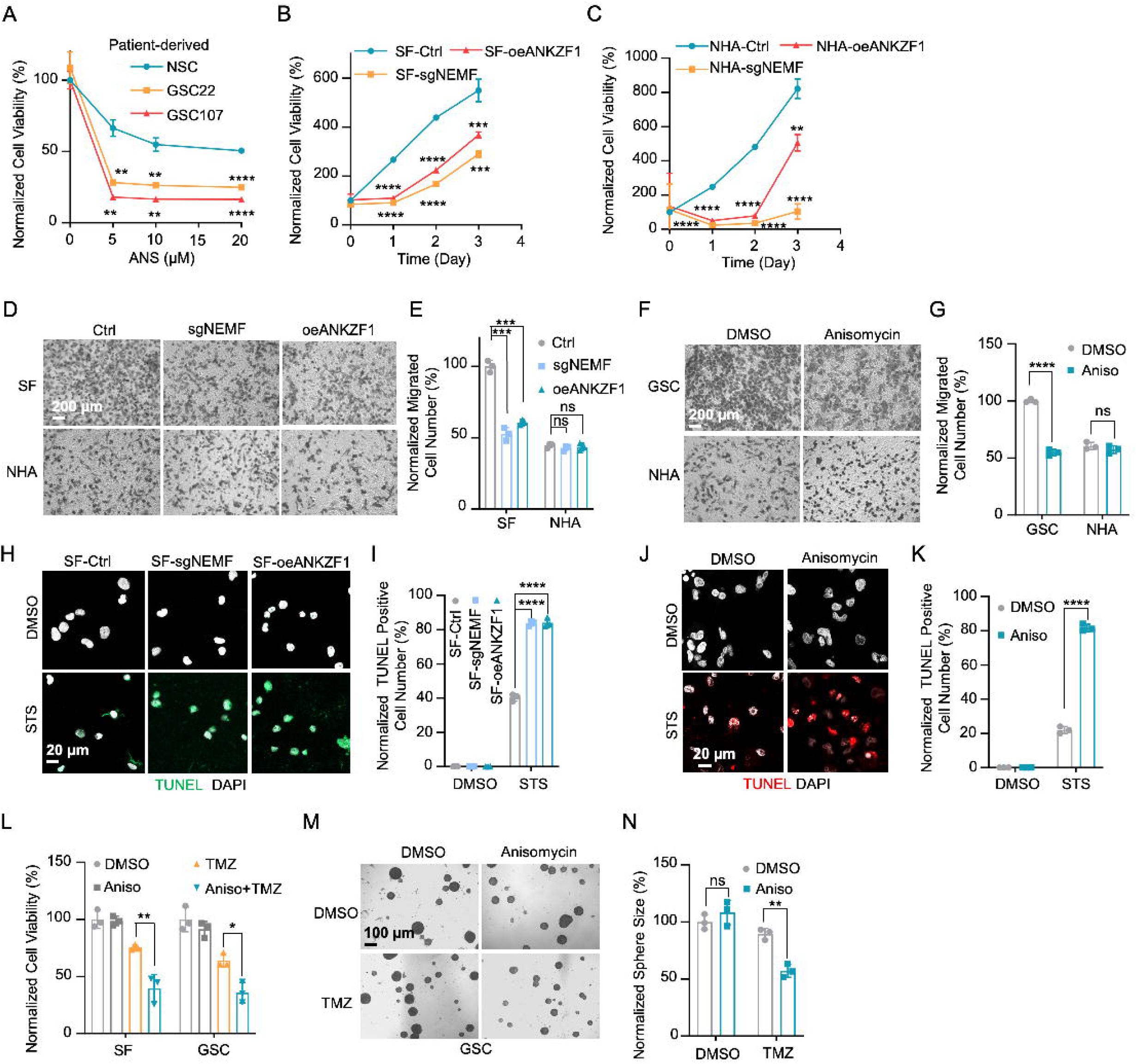
Inhibition of msiCAT-tailing impedes GBM progression. (A) Cell viability assay shows greater sensitivity to anisomycin treatment in patient-derived GSC cells than control NSC cells at 48 hours (n=3; unpaired Student’s t-test; **, *P* < 0.001; ****, *P* <0.0001; compared to controls at the corresponding dose). (B) MTT assay indicates reduced GBM cell proliferation by genetic inhibition of the msiCAT-tailing pathway (n=3; unpaired Student’s t-test; **, *P* < 0.01; ***, *P* < 0.001; ****, *P* <0.0001, compared to controls at the corresponding time). (C) MTT assay indicates reduced NHA cell proliferation by genetic inhibition of the msiCAT-tailing pathway (n=3; unpaired Student’s t-test; **, *P* < 0.01; ****, *P* <0.0001, compared to controls at the corresponding time). (D, F) Transwell assay reveals that both genetic (D) and pharmacological (F) inhibition of the msiCAT-tailing pathway hampers the migration of GBM cells but not control cells. (E, G) Quantification of (D, F) showing the number of migrated cells (n=3; unpaired Student’s t-test; ***, *P* < 0.001; ****, *P* <0.0001; ns, not significant). (H, J) TUNEL staining reveals that both genetic (H) and pharmacological (J) inhibition of the msiCAT-tailing pathway promote STS-induced apoptosis in GBM cells, utilizing TUNEL-Cy3 as an apoptotic cell marker and DAPI as a nuclear stain. (I, K) Quantification of (H, J) showing the percentage of TUNEL-positive cells in the population (n=3; unpaired Student’s t-test; **** *P* < 0.0001), using DMSO as the vehicle control. (L) MTT assay shows that pharmacological inhibition of the msiCAT-tailing pathway decrease the resistance of GBM cells to temozolomide (TMZ, 150 µM) treatment (n=3; unpaired Student’s t-test; * *P* < 0.05; **, *P* < 0.01). (M) The neurosphere formation assay shows that reduced spheroid formation, caused by pharmacological inhibition of the msiCAT-tailing pathway, can synergize with TMZ in GBM cells. (N) Quantification of (M) (n=3; unpaired Student’s t-test; **, *P* < 0.01).

Further investigation revealed the RQC pathway’s involvement in GBM cell anti-apoptosis, with initial findings pointing to alterations in mitochondrial functions. Prior studies demonstrated that genetic or pharmacological inhibition of the RQC pathway led to a significant decrease in GBM mitochondrial membrane potential (ΔΨm) (Figure 2C, D). In GSC cells, anisomycin treatment promoted mitochondrial permeability transition pore (MPTP) opening, an effect not seen in NHA cells (Figure 3A-D). Consequently, GBM cell lines with genetically or pharmacologically inhibited RQC pathways were more susceptible to STS-induced apoptosis, evidenced by elevated executioner caspase 3/7 activity (Figure 5 – Figure Supplement 1A), enhanced PARP-1 cleavage (Figure 5 – Figure Supplement 1B, C), increased TUNEL staining (Figure 5H-K), and flow cytometry analysis (Figure 5 – Figure Supplement 1D-G). Notably, general translation inhibition using cycloheximide did not elicit the same apoptotic response (Figure 5 – Figure Supplement 1A, D, E). Finally, the RQC pathway was also implicated in temozolomide (TMZ)-induced cell death. Combining anisomycin with TMZ significantly reduced GBM cell survival (Figure 5L) and effectively inhibited GSC spheroid growth (Figure 5M, N). In summary, the RQC pathway plays a critical role in multiple aspects of GBM progression, including proliferation, migration, and survival under apoptotic stress.

## Discussion

The Ribosome-associated Quality Control (RQC) pathway plays a crucial role in managing aberrant proteins produced during translation. This study focused on understanding the consequences of RQC-mediated modification, specifically the addition of msiCAT tails, on mitochondrial proteins such as ATP5α in glioblastoma (GBM) cells. The findings reveal that GBM cells harboring msiCAT-modified ATP5α exhibit a unique metabolic profile. Despite a reduction in ATP synthesis, these cells maintain their mitochondrial membrane potential (ΔΨm), a key factor for cellular function and survival. Furthermore, they demonstrate enhanced cell survival and motility, characteristics associated with increased tumor invasiveness and metastasis. Notably, the presence of msiCAT-modified ATP5α confers resistance to apoptosis triggered by staurosporine (STS), potentially by modulating the mitochondrial permeability transition pore (MPTP), a critical regulator of cell death pathways, as illustrated in Figure 6. These identified traits contribute to an increased aggressiveness of tumors, suggesting that the RQC pathway plays a critical role in cancer cell survival and proliferation. Encouragingly, a recent study also demonstrated the RQC pathway’s involvement in a *Drosophila* model of Notch overexpression-induced brain tumors (56). The findings imply modulating the RQC pathway could serve as a promising complementary strategy to existing chemotherapy regimens. By targeting this specific pathway, therapeutic interventions might effectively disrupt the mechanisms that allow cancer cells to evade apoptosis and sustain their energy production under stress, potentially leading to improved treatment outcomes for patients with GBM and other cancers characterized by similar protein modifications.

**Figure 6.**
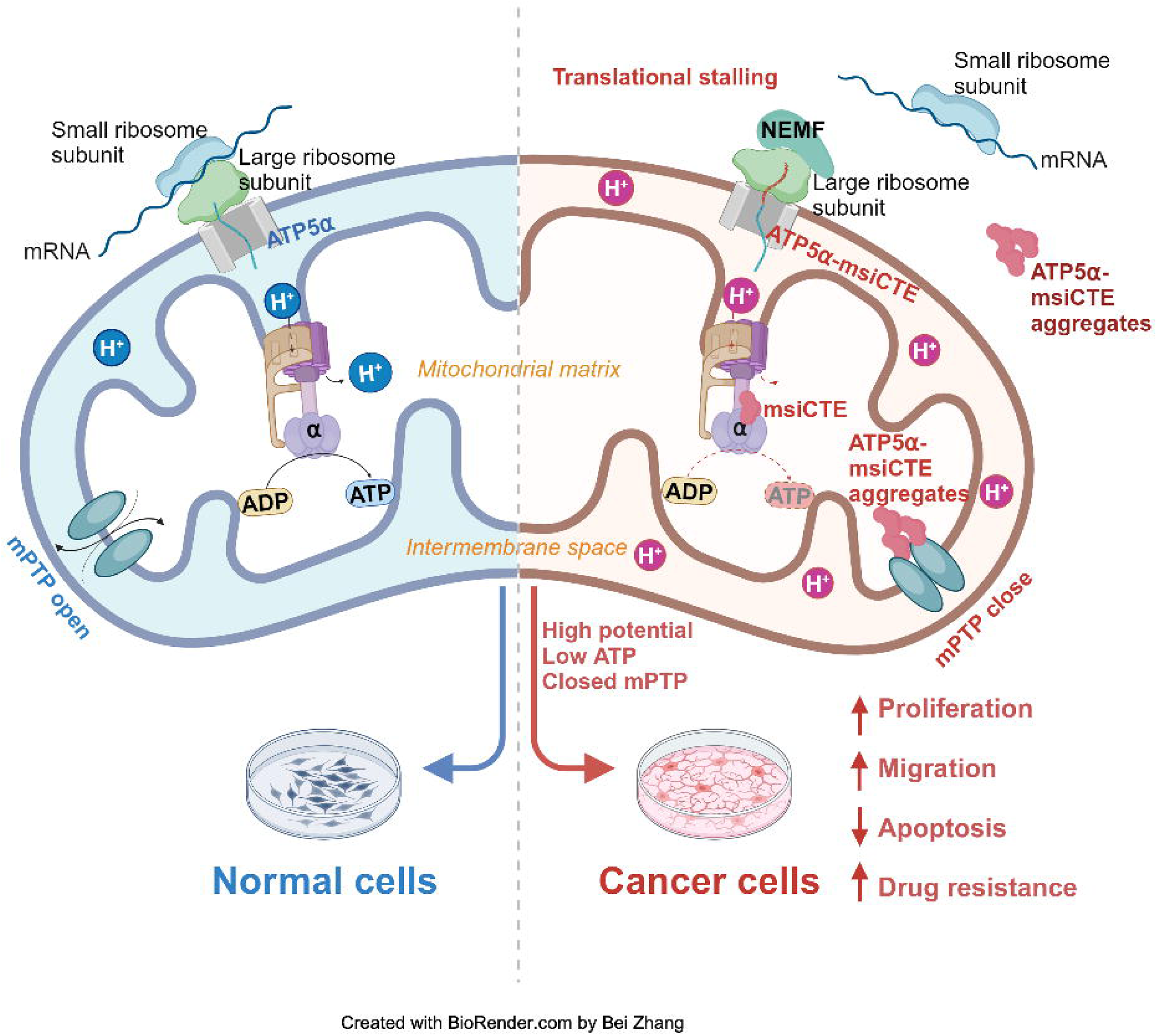
Impact of msiCAT-tail modified ATP5α protein on mitochondrial function in GBM cells. In healthy cells, ATP5α protein, encoded by the nuclear genome, is imported into the mitochondrial matrix via the TOM/TIM complex through co-translational import and incorporated into ATP synthase (Left). Conversely, in GBM cells, the CAT-tailed ATP5α protein can either form aggregates near the mitochondrial outer membrane or be imported into the mitochondria. Within the mitochondrial matrix, proteins with shorter CAT-tails readily integrate into ATP synthase, disrupting its functionality. This dysfunction is characterized by a reduced ATP synthesis rate and proton (H^+^) accumulation, resulting in an elevated mitochondrial membrane potential (ΔΨm). These alterations in ATP synthase ultimately trigger malfunction of the MPTP, consequently affecting cell proliferation, migration, and resistance to drug-induced apoptosis (Right).

The study of ATP synthase behavior in cancer holds particular importance. During carcinogenesis, ATP synthase frequently relocates to the plasma membrane, where it is referred to as ectopic ATP synthase (eATP synthase). These eATP synthases exhibit catalytic activity, facilitating ATP production in the extracellular space to foster a favorable tumor microenvironment (57). Research indicates that eATP synthase assembles initially in mitochondria before being transported to the cell surface via microtubules (47). However, the specific type of ATP synthase delivered to the plasma membrane remains unclear. Future investigations into the localization of CAT-tailed eATP synthase may offer valuable insights into this process.

Multiple mitochondrial proteins in cancer cells can likely undergo CAT-tailing in a similar way. These msiCAT-tailed peptides may have varied impacts on mitochondria and cells due to differences in their base proteins. For instance, CAT-tailed COX4 protein might substantially and directly diminish mitochondrial respiratory efficiency. Examining the individual roles of these proteins is important, as the combined effect of their defects may be crucial in understanding observed mitochondrial changes in cancer. A minor caveat here is that the observed effect of the CAT-tails’ presence primarily stems from artificial CAT-tail sequences with a high threonine content, rather than the endogenous CAT-tail protein. It is possible that other sequence components could lead to different effects (44). A recent study found that ANKZF1 knockdown inhibited GBM progression by causing abnormal protein accumulation in mitochondria (29). This, combined with our data, suggests that balanced ANKZF1 expression and activity are vital for cancer proliferation. Both excess and deficiency may alter cellular adaptability. A minor flaw of that study was the use of a mitochondrial-localized non-stopped GFP protein to induce proteostasis stress and the lack of direct biochemical evidence of CAT-tailed proteins. Our research focuses on endogenous proteins for a detailed analysis of their impact on mitochondria. The rationale is that highly expressed, non-physiological ectopic proteins might cause general proteostasis failure, masking the specific functions of endogenous proteins. Additionally, the studies used different cell lines. GSC, a patient-derived GBM cell line with greater stemness, might have a distinct mitochondrial status and RQC pathway activity compared to U87 or U251 cell lines. Thus, the conclusions of the two studies are not contradictory but rather complementary, both demonstrating the significance of RQC in tumorigenesis. Our study delves into the mechanistic role of the RQC pathway in GBM, identifying new potential targets for future treatments.

An in-depth investigation into the quantification of nuclear genome-encoded mitochondrial proteins modified via the msiCAT-tailing mechanism using sophisticated mass spectrometry is a compelling area for future research. Recent work by Lv et al., published in *Cell Reports*, revealed that the cytoplasmic E3 ligase Pirh2 and the mitochondrial protease ClpXP work in conjunction with the established NEMF-ANKZF1 system to break down mitochondrial protein aggregates resulting from ribosome stalling (58). The increased presence of ClpXP in various cancers could potentially be linked to an increase in msiCAT-tailing products in mitochondria, though further studies are needed to clarify ClpXP’s role in mitochondrial RQC (59). Moreover, ClpXP influences the levels of multiple mitochondrial proteins. Our own experiments showed that ATP5α proteins lacking msiCAT-tails were the most challenging to express ectopically. Proteins with shorter tails (AT3) expressed more readily, while those with longer tails (AT20) exhibited the highest expression levels but also tended to form SDS-insoluble aggregates. This regulatory effect could be mediated by ClpXP-dependent degradation or potentially through transcriptional control. PGC-1α, the peroxisome proliferator-activated receptor gamma co-activator, is a key regulator of mitochondrial biogenesis in mammals (60). By binding to and activating nuclear transcription factors, PGC-1α triggers the transcription of nuclear genome-encoded mitochondrial proteins and the mitochondrial transcription factor Tfam. Tfam, in turn, activates mitochondrial genome transcription and replication (61). Distinguishing between these regulatory possibilities will necessitate future research, including a meticulous examination of mRNA levels for msiCAT-tailed targets and analysis of PGC-1α and Tfam binding to transcriptional elements.

MPTP is a complex, supramolecular channel traversing the inner mitochondrial membrane, characterized by its non-selective ion permeability, calcium dependence, and multifaceted functionality. Despite extensive investigations into its functional attributes and regulatory mechanisms, the precise molecular architecture of the MPTP remains elusive (62). Several theoretical models have been posited to elucidate the MPTP’s structural composition. Firstly, the VDAC/ANT/Cyp-D model (63) proposed an assembly of voltage-dependent anion channels (VDAC), adenine nucleotide translocators (ANT), and cyclophilin D (Cyp-D) as the structural basis; however, subsequent genetic analyses have introduced substantial controversy regarding the integral role of these proteins within the MPTP complex (64–67). Secondly, the ATP synthase model posits that MPTP formation involves dimers or reconstituted c-rings of ATP synthase (48, 49). While this hypothesis presents an intriguing perspective, empirical confirmation of ATP synthase’s role as a definitive structural element of the pore remains inconclusive, with a body of conflicting research surrounding this proposition. Thirdly, the contemporary prevailing hypothesis suggests the MPTP is constituted by a large complex, termed the ATP synthasome, comprising ANT and ATP synthase, with Cyp-D serving a regulatory function over the complex’s dynamic behavior (68).

The MPTP activity is modulated by mitochondrial membrane potential (ΔΨm), which reciprocally influences mitochondrial ion homeostasis and energy metabolism (69, 70). Our study elucidates a dual function of msiCAT-tailed ATP5α protein in cancer cells: stabilization of a high membrane potential, thereby mitigating MPTP induction, and direct inhibition of MPTP functionality through participation in its assembly. While MPTP’s critical role in cell death is established, the premise that MPTP inhibition enables cancer cell evasion of drug-induced programmed cell death has lacked substantial evidence. This study furnishes empirical support for this hypothesis, demonstrating that GBM cells, notably glioblastoma stem cells (GSC), exhibit markedly reduced MPTP activity relative to control cells. This reduced activity is directly correlated with the CAT-tailing modification of the ATP synthase subunit. These observations are concordant with prior research indicating that genetic mutations or post-translational modifications in specific ATP synthase subunits can modulate MPTP activity. The findings highlight a novel mechanism through which cancer cells may develop resistance to therapeutic interventions by manipulating mitochondrial function (71, 72).

## Materials and Methods

### Key resources table

Please see the appendix.

### Cell lines and cell culture conditions

The human astroglia cell line SVG p12 (ATCC, cat. CRL-8621) and the human glioma cell line SF268 were from Dr. Rongze Olivia Lu. Both cell lines were cultured in DMEM (ATCC, cat. #302002) with 10% FBS (Biowest, cat. S1620-100) and penicillin/streptomycin (Gibco™, cat. 15140122). SF268 clones should be maintained in complete DMEM supplemented with 400 µg/mL G418 (Gibco, cat. 10131027). The 0.25% trypsin solution (ATCC, cat. #SM2003C) was used to passage cells. The normal human astrocytes NHA E6/E7/hTERT cell line was from Dr. Russell O. Pieper, UCSF Brain Tumor Research Center. Cells are cultured in ABM^TM^ Basal Medium (Lonza, cat. CC-3187) and AGM^TM^ SingleQuots^TM^ Supplements (Lonza, cat. CC-4123). Corning™ Accutase™ Cell Detachment Solution (Corning, cat. 25058CI) was used to passage cells. GSC827, a patient-derived human glioma stem cell line, was from Dr. Chun-Zhang Yang at NIH. The NSC, NSC26, patient-derived GSC33, GSC22, GSC99, GSC105, and GSC107 cell lines used in this study were kindly provided by Dr. John S Kuo at the University of Texas, Austin. Derivation of these lines from patient GBM specimens is described earlier (73). Detailed characterizations of the GSC lines (not GSC105 & 107) are available in their previous publication (74). GSC 105 & 107 are not previously published. GSC cells were cultured in Neural basal-A Medium (Gibco, cat. #10888022) with 2% B27 (Gibco, cat. #17504044), 1% N2 (Gibco, cat. #17502048), 20 ng/ml of EGF and FGF (Shenandoah Biotechnology Inc. cat. PB-500-017), Antibiotic-Antimycotic (Gibco, cat. #15240062), and L-Glutamine (Gibco, cat. #250300810). Cells could be cultured in both spherical and attached (on Geltrex, Thermo Fisher, cat. A1413202) forms. Corning™ Accutase™ Cell Detachment Solution (Corning, cat. 25058CI) was used to passage cells.

Cells were transfected with X-tremeGENE™ HP DNA Transfection Reagent (Sigma, cat. 6366244001) following the standard protocol. For single clone selection, SF268 cells were treated with 800 µg/ml G418 for 5 days. The cells were then seeded into a 96-well plate at a density of 1/100 µL. Positive clones were verified by immunofluorescence staining and immunoblotting. Cells were maintained in complete DMEM containing 400 µg/mL G418. GBM cell lines were subjected to a 4-hour pre-treatment at 37°C using either anisomycin (20 nM or 200 nM, Fisher Scientific, cat. AAJ62964MF) or cycloheximide (100 µg/mL, Fisher Scientific, cat. AC357420010) in medium, as detailed in the conducted experiments.

### Primers, plasmids, and viruses

Plasmids pcDNA3.1+/C-(K)-DYK-ATP5F1A (pATP517 control), pcDNA3.1+/C-(K)-DYK-ATP5F1A-AT3 (pATP517-AT3), pcDNA3.1+/C-(K)-DYK-ATP5F1A-AT20 (pATP517-AT20), pcDNA3.1+/C-(K)-DYK-ATP5F1A-GS3 (pATP517-GS3), and pcDNA3.1+/C-(K)-ATP5F1A-DYK-GS20 (pATP517-GS20) were generated by GenScript Inc. Plasmids pCMV-5×FLAG-β-globin-control (5FBG-Ctrl) and pCMV-5×FLAG-β-globin-non-stop (5FBG-nonstop) were generated by Dr. Hoshino (Nagoya City University) and Dr. Inada (Tohoku University) (43). pCMV6-DDK-NEMF (oeNEMF) was from ORIGENE Inc. (cat. RC216806L3).

Viruses (and plasmid used to generate viruses) are pLV[CRISPR]-hCas9:T2A:Neo-U6>Scramble[gRNA#1] (sgControl/sgCtrl), pLV[CRISPR]-hCas9:T2A:Neo-U6>hNEMF[gRNA#1579] (sgNEMF), pLV[Exp]-Bsd-EF1A>ORF_Stuffer (pLV-control), pLV[Exp]-EGFP:T2A:Puro-EF1A>mCherry (pLV-control-2/oeCtrl), pLV[Exp]-Bsd-EF1A>hANKZF1[NM_001042410.2]/HA (oeANKZF1), and pLV[Exp]-mCherry/Neo-EF1A>hANKZF1[NM_001042410.2] (oeANKZF1) were made by VectorBuilder Inc.

Primers (5’ to 3’) used for RT-PCR are:

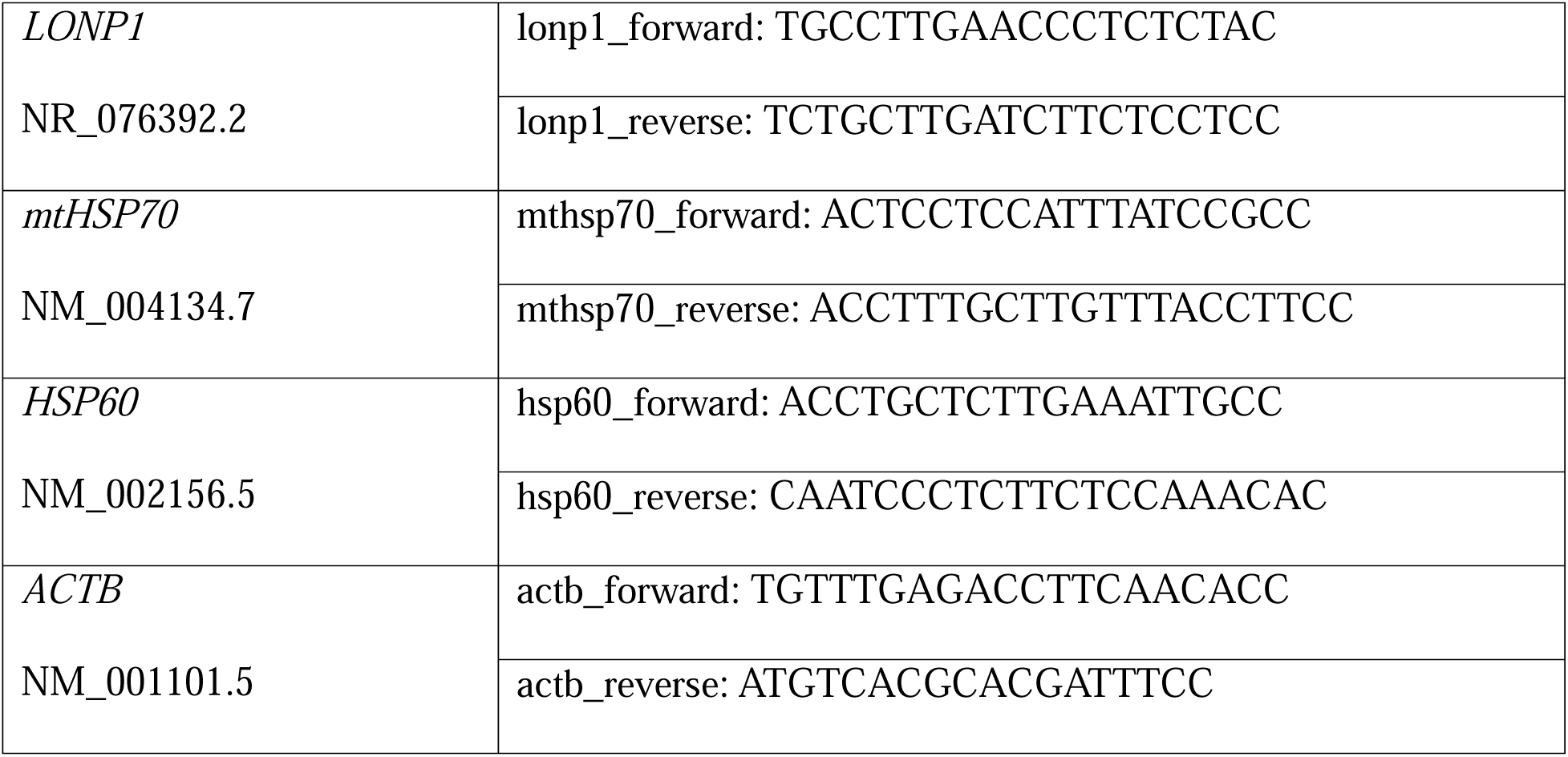

### Neurosphere formation assay of GSCs

The GSC spheroids were dissociated using Accutase for 2 min. Cells were resuspended in a single-cell suspension and grown under non-adherent conditions. Cells were seeded in 12-well plates at a density of 0.25 x 10^6^ cells/well and cultured in 3 mL culture medium for 24 hours. 20 nM of anisomycin and 150 µM of temozolomide (TMZ) were added to the culture medium and the cells were treated for 96 hours. Spheroids were imaged under a 10x objective, captured using QCapture, and analyzed with ImageJ. Spheroids larger than 50 µm were counted.

### Differential gene expression analysis using the public database

The raw RNA-seq data used for the analysis were obtained from the University of California, Santa Cruz Xenabrowser (cohort: TCGA TARGET GTEx, dataset ID: TcgaTargetGtex_rsem_gene_tpm, https://xena.ucsc.edu/). Subsets were then created to include only TCGA glioma (GBM), GTEx Brain Frontal Cortex, and GTEx Cortex samples. Differential expression analysis was conducted using the “Limma” package (R version: 4.3.1). The Benjamini-Hochberg method was used for multiple testing correction to control the false discovery rate (FDR). Cut-off of adjusted p-value (adj.P.Val) was set at 0.001; cut-off of the absolute fold change was set at 2 (logFC > 1). The code is available without restrictions at https://github.com/yuanna23/GBM_elife.

### Immunostaining

Cells were cultured on sterile coverslips until 80% confluency. For immunostaining, cells were washed with phosphate-buffered saline (PBS) solution thrice. Then, 4% formaldehyde (Thermo Fisher, cat. BP531-500) was applied to cells for fixation for 30 min at room temperature. After fixation, cells were washed with PBS solution containing 0.25% Triton X-100 (PBSTx) (Thermo Fisher, cat. T9284) thrice, and blocked with 5% normal goat serum (Jackson Immuno, cat. 005-000-121) for 1 hour at room temperature. Cells were then incubated with primary antibodies overnight in a humidified chamber at 4°C. The next day, cells were washed by PBSTx thrice and incubated with secondary antibodies for 2 hours at room temperature. After washing, cells were stained with 300 nM DAPI (Thermo Fisher, cat. 57-481-0) for 5 min at room temperature and mounted in Fluoromount-G Anti-Fade solution (Southern Biotech, cat. 0100-35). Images were captured using a Zeiss LSM 800 confocal microscope with a 40x oil objective lens and AiryScan processing. The primary antibodies used in the study were rabbit anti-ATP5a (Cell Signaling, cat. #18023), mouse anti-TOMM20 (1:500, Santa Cruz, cat. sc-17764), rabbit anti-MTCO2 (1:500, Proteintech, cat. 55070-1-AP), and mouse anti-NDUS3 (1:1000, Abcam, cat. ab14711). The secondary antibodies were Alexa fluor 633-, 594-, 488-conjugated secondary antibodies (1:300, Invitrogen, cat. A21071, A11036, A32732).

### SDS-PAGE and immunoblotting

Cells or isolated mitochondria were solubilized in cell lysis buffer containing 50 mM Tris-HCl pH 7.4, 150 mM NaCl, 10% glycerol, 1% Triton X-100, 5 mM EDTA, and 1x protease inhibitor (Bimake, cat. B14002). Protein concentration was measured by using the Bradford assay (BioVision, cat. K813-5000-1). Samples were separated in a 4-12% Tris-Glycine gel (Invitrogen, cat. WXP41220BOX) and proteins were transferred to a PVDF membrane (Millipore, cat. ISEQ00010). The membranes were then blocked with 5% non-fat dry milk (Kroger) for 50 min at room temperature and probed with primary antibodies overnight at 4°C. Membranes were washed with Tris-buffered saline with 0.1% Tween 20 (TBST) solution thrice and then incubated with secondary antibodies for 1 hour at room temperature. Blots were detected with ECL solution (PerkinElmer, cat. NEL122001EA) and imaged by Chemidoc system (BioRad). The intensity of blots was further analyzed by ImageJ software. The primary antibodies used were mouse anti-Actin (1:1000, Santa Cruz, cat. sc-47778), rabbit anti-NEMF (1:1000, Proteintech, cat. 11840-1-AP), mouse anti-ANKZF1 (1:1000, Santa Cruz, cat. sc-398713), mouse anti-ATP5a (Abcam, cat. ab14748), mouse anti-NDUS3 (1:1000, Abcam, cat. ab14711), rabbit anti-COX4 (Abcam, cat. ab209727), mouse anti-Flag (1:1000, Sigma, cat. F1804), rabbit anti-ANT1/2 (1:1000, Proteintech, cat. 17796-1-AP), rabbit anti-CypD (1:1000, Proteinetch, cat. 15997-1-AP), rabbit anti-PARP1 (1:1000, Abclonal, cat. A0942), rabbit anti-GAPDH (1:1000, Abclonal, cat. A19056). The secondary antibodies used were goat anti-rabbit IgG (1:5000, Invitrogen, cat. G21234), goat anti-mouse IgG (1:5000, Invitrogen, cat. PI31430).

### Mitochondrial isolation, blue Native PAGE, and western blotting

Cells were homogenized using Dounce homogenizer in ice-cold homogenization buffer containing 210 mM mannitol (Fisher Sci, cat. AA3334236), 70 mM sucrose (Fisher Sci, cat. AA36508A1), 5 mM HEPES (Fisher Sci, cat. 15630106), pH 7.12, 1 mM EGTA (Fisher Sci, cat. 28-071-G), and 1x protease inhibitor. The homogenate was centrifuged at 1500 g for 5 min. The resultant supernatant was centrifuged at 13000 g for 17 min. The supernatant was collected as the cytosol portion, and the pellet (the mitochondria portion) was washed with homogenization buffer and centrifuged at 13000 g for 10 min. For Blue Native PAGE, the mitochondrial samples were solubilized in 5% digitonin (Thermo Fisher, cat. BN2006) on ice for 30 min and then centrifuged at 20,000 g for 30 min. The supernatant contains solubilized mitochondrial proteins and was mixed with 5% G-250 (GoldBio, cat. C-460-5) and 1x NativePAGE sample buffer (Invitrogen, cat. BN2008) (final G-250 concentration is 25% of the digitonin concentration). Mitochondrial protein concentration was measured by using the Bradford assay. Samples were separated in 3-12% Bis-Tris Native gel (Invitrogen, cat. BN1001BOX) and then transferred to a PVDF membrane. Membranes were fixed with 8% acetic acid (Thermo Fisher, cat. 9526-33), and then blocked and probed with antibodies as described above for Western blotting.

### Mitochondrial membrane potential assays

Mitochondrial membrane potential of GSC cells was measured using Image-iT^TM^ TMRM (Invitrogen, cat. I34361). Cells were cultured in 96-well black plates at a density of 1 x 10^5^ cells per well overnight in an incubator with 5% CO_2_ at 37°C. Cells were incubated with TMRM (100 nM) for 30 min at 37°C. Then, cells were washed with PBS solution three times. Fluorescence changes at excitation/emission of 548/574 nm were monitored with a Cytation 5 plate reader (BioTek). Mitochondrial membrane potential was also measured using JC-10 (AdipoGen, cat. 50-114-6552). Cells were cultured in 96-well black plates at a density of 5 x 10^4^ cells per well overnight in an incubator with 5% CO_2_ at 37°C. Cells were incubated with JC-10 (10 µg/ml) for 45 min at 37°C. Then, cells were washed with PBS solution twice. Fluorescence changes at excitation/emission of 535/595 nm for JC-10 aggregates and at 485/535 nm for JC-10 monomers were monitored with a Synergy 2 Reader (BioTek). Mitochondrial membrane potential was quantified as the fluorescence of JC-10 aggregates/monomers (595/535 nm).

### Seahorse cell mitochondrial stress assays

The oxygen consumption rate (OCR) of cells was measured using the Seahorse Cell Mito Stress Test kit following the user guide (Agilent, cat. 103010-100). Briefly, cells were cultured overnight in testing chambers at a density of 8,000 cells per well in an incubator with 5% CO_2_ at 37°C. Cells were then washed twice with assay medium containing Seahorse XF DMEM medium (Agilent, cat. 103575-100) supplemented with 1 mM pyruvate, 2 mM glutamine, and 10 mM glucose. They were subsequently incubated in the assay medium for 1 hour in an incubator without CO_2_ at 37°C. Cells were treated with compounds in the order of oligomycin (1.5 µM), carbonyl cyanide-4 (trifluoromethoxy), phenylhydrazone (FCCP, 1.0 µM), and Rotenone/Antimycin (0.5 µM). The OCR of cells was monitored by using Seahorse XF HS Mini (Agilent).

### Mitochondrial MPTP assay

The status of mitochondrial permeability transition pore was measured using Invitrogen™ Image-IT™ LIVE Mitochondrial Transition Pore Assay Kit (Invitrogen, cat. I35103). Cells were cultured in 35 mm glass-bottom dishes overnight in an incubator with 5% CO_2_ at 37°C. Cells were washed twice with the modified Hank’s Balanced Salt Solution (HBSS, Thermo Fisher, cat. 14025092) containing 10 mM HEPES, 2 mM L-glutamine and 0.1 mM succinate (Thermo Fisher, cat. 041983.A7) and incubated with the labeling solution (1 µM Calcein, 0.2 µM MitoTracker Red, 1 mM Cobalt Chloride) for 15 min at 37°C. Cells were then washed with HBSS twice and imaged at excitation/emission of 494/517 nm for Calcein and at 579/599 nm for MitoTracker Red by using the Zeiss confocal microscope.

### Mitochondrial Ca^2+^ retention capacity assay

The mitochondrial calcium retention capacity (CRC) was measured on a Cytation 5 reader at excitation/emission of 506/592 nm using the membrane-impermeable fluorescent probe Calcium green-5N (Invitrogen, cat. C3737). Isolated mitochondria samples (0.75 mg protein/mL) were incubated in 1 mL swelling medium supplemented with 10 mM succinate, 1 μM Calcium green-5N, inorganic phosphate, and cyclosporine A (Thermo Fisher, cat. AC457970010). One Ca^2+^ addition was 1.25 nmol (1 mL volume). Only the MPTP opening in the presence of cyclosporine A was induced by high amounts of added calcium (30 nmol Ca^2+^ in the last two additions). The CRC value was calculated as total Ca^2+^ accumulated in the mitochondria per unit (1 mg protein).

### MTT assay

Cell proliferation was measured by using the MTT assay kit (Roche, cat. 11465007001). Cells were cultured in 96-well plates at a density of 2000 cells per well overnight in an incubator with 5% CO_2_ at 37°C. Cells were treated with MTT labeling reagent for 4 hours at 37°C. The solubilization buffer was added to the cells, and then the cells were incubated overnight at 37°C. Absorbance changes of the samples at 550 nm were monitored by using a Synergy 2 Reader (BioTek).

### Wound healing assay

Cells were seeded into 6-well plates and cultured for 24-48 hours to reach a confluent cell monolayer. Cells were treated with serum-free medium overnight before mechanical scratching (54). Images of the wounds were taken at 0, 24, and 48 hours. Wound areas were measured by using the wound healing plugin of ImageJ. Wound Coverage % = 100% x [A*_t=0h_*-A*_t=Δh_*]/A*_t=0h_*(A*_t=0h_* is the area of the wound measured immediately after scratching *t = 0h*, A*_t=Δh_* is the area of the wound measured *h* hours after the scratch is performed).

### Cell migration assay

Cell migration was measured by using Transwell assays (Corning, cat. CLS3422). Cells were cultured in Transwell inserts at a density of 1 x 10^5^ cells per well for 3 hours in an incubator at 37°C with 5% CO_2_. The top inserts were supplemented with DMEM medium only, and the bottom wells were supplemented with DMEM medium with 20% Fetal Bovine Serum. After incubation, the cells on the apical side of the Transwell insert membrane were removed using a cotton applicator. The cells on the bottom side of the insert were rinsed with PBS twice and fixed in 70% ethanol (Thermo Fisher, cat. R40135) for 15 min at room temperature. After fixation, inserts were placed into an empty well to allow the membrane to dry. Then, the insert was incubated with 0.2% crystal violet (Sigma, cat. V5265) for 5 min at room temperature. The insert was rinsed with water twice, and images were captured by using a microscope with a 20x objective. Cell numbers were quantified using ImageJ.

### TUNEL staining

The apoptosis was measured by a TUNEL assay kit (ApexBio, cat. K1134). Cells were cultured on sterile cover slips until 80% confluency and washed with PBS thrice. Then, 4% formaldehyde was applied to cells and fixed for at 4°C 25 min. After fixation, cells were washed with PBS twice and incubated with 20 µM proteinase K (Invitrogen, cat. 25530049) for 5 min at room temperature. Then, cells were rinsed with PBS thrice and incubated in 1x equilibration buffer for 10 min at room temperature. Cells were stained with FITC or Cy3 labeling mix for 1 hour at 37°C in a humidified chamber. Cells were washed by PBS thrice and stained with DAPI for 5 min at room temperature. Cells were mounted in the Fluoromount-G Anti-Fade solution and imaged at 520 nm for FITC or at 570 nm for Cy3 by using the Zeiss confocal microscope.

### Caspase-3/7 activity assay

Caspase-3/7 activity was measured by using CellEvent™ Caspase 3/7 Detection Reagents (Invitrogen, cat. C10432) following the manufacturer’s protocol. Specifically, cells were seeded in a 96-well black plate with a clear bottom at a density of 5 x 10^4^ cells per well and incubated overnight in the incubator with 5% CO_2_ at 37°C. Cells were then incubated with 1x staining solution for 30 min at 37°C. Fluorescence changes at excitation/emission of 485/525 nm were monitored with a Synergy 2 Reader (BioTek).

### Annexin V-FITC/Propidium Iodide (PI) apoptosis detection

Annexin V-FITC/PI apoptosis assay was performed by using the FITC Annexin V Apoptosis Detection Kit with PI (BioLegend, cat. 640914). Briefly, 1 x 10^5^ cells were collected in 100 μL of staining buffer. Then, cells were incubated with 5 μL of Annexin V-FITC and 2.5 μL of PI for 15 min at room temperature in the dark. Following incubation, 400 μL of binding buffer was added to the stained cells. Flow cytometry analysis of the fluorescence was performed using a Soni SH800 Cell Sorter.

### Mitochondria ATP measurement via fluorescence imaging of ATP-red

BioTracker™ ATP red dye (Millipore, cat. SCT045) is a fluorogenic indicator for ATP in mitochondria (75). Cells cultured in monolayer conditions were incubated in medium with 5 μM ATP red for 15 min in an incubator at 37°C with 5% CO_2_. Mitochondria were also labeled by incubating cells with 100 nM MitoTracker-Green (Invitrogen, cat. M7514) for 15 min to normalize their mass. Before measurement, cells were washed twice with culture medium, and then fresh medium was added. Cells were imaged in a 37°C chamber with 5% CO_2_ at excitation/emission of 510/570 nm for ATP-red and at excitation/emission of 490/516 nm for MitoTracker-Green by using the Zeiss confocal microscope. The ATP-red signals could also be measured by a Synergy 2 Reader (BioTek).

### Co-immunoprecipitation

Cells were lysed in the buffer containing 50 mM Tris-HCl pH 7.4, 150 mM NaCl, 10% glycerol, 1% TritonX-100, 5 mM EDTA, and 1x protease inhibitor. Soluble samples were incubated with 1.5 µL ATP517 antibody at 4°C with mixing overnight. 25 µL of protein A/G magnetic beads (Pierce, cat. 88802) were added to the co-IP samples and incubated at 4°C with mixing overnight. Samples were washed with washing buffer thrice and then applied to SDS-PAGE analysis.

### Mice and immunostaining

Animal studies were approved by the University of California, San Francisco Institutional Animal Care and Use Committee (IACUC, AN195636-01) and were performed following the guidelines of the National Institutes of Health (NIH).

For orthotopic brain tumor models, 8-to-10-week-old C57BL/6J mice (male and female in equal numbers) were used for i.c. studies. Cell lines (GL261, SB28) were suspended in DMEM for inoculation. Mice were anesthetized with isoflurane, and 30,000 tumor cells were injected orthotopically in 3 μL. Using a stereotactic frame, a burr hole was formed on the skull via a 0.7 mm drill bit 1.5mm laterally to the right and 1.5mm rostrally from the bregma, and a noncoring needle (26s gauge; Hamilton) was used to inject the cells at a depth of 3mm into the brain from the burr hole. The skin incision was sutured. Mice were then monitored daily. Mouse SB28 tumor tissue and wild-type mouse brain tissue were collected at the survival endpoint.

Frozen tissue sections were thawed at room temperature for 20 min and rinsed with PBS three times. Tissues were then fixed in 4% formaldehyde for 15 min at room temperature. After washing in PBS, tissues were permeabilized with 0.01% Triton X-100 + 0.1% Tween-20 for 15 min and then blocked by using 5% normal goat serum and M.O.M. blocking reagent (Vector Laboratories, cat. BMK-2202) for 1 hour at room temperature. Tissues were then incubated with primary antibodies overnight in a humidified chamber at 4°C. After washing in PBST, tissues were incubated with secondary antibodies for 1 hour. After washing again in PBST, tissues were stained with 300 nM DAPI for 5 min and mounted in Fluoromount-G Anti-Fade solution. Images were taken using a Zeiss LSM 800 confocal microscope. The primary antibodies used in the study were mouse anti-ATP5a (1:500, Abcam, cat. Ab14748), rat anti-TOMM20 (1:500, Abcam, cat. Ab289670), rabbit anti-NEMF (1:500, Proteintech, cat. 11840-1-AP), mouse anti-ANKZF1 (1:500, Santa Cruz, cat. sc-398713), chicken anti-GFP (1:500, Abcam, cat. Ab13970). The secondary antibodies were Alexa Fluor 633-, 594-, 488-conjugated secondary antibodies (1:300, Invitrogen, cat. A21071, A11036, A32732).

## Statistics

Statistical analyses were performed using GraphPad Prism 9.4. Chi-squared test and unpaired Student’s t-test were used for comparison. *P* < 0.05 was considered significant, except in gene expression analysis (Figure. 1A). *, *P* < 0.05; **, *P* < 0.01; ***, *P* < 0.001; ****, *P* < 0.0001; ns, not significant. All data were expressed as means ± s.e.m. This study’s replicates, samples, groups, and experiments were biologically independent, except in Table 1. The “n” numbers for each assay are indicated in the figure legends.

## Materials availability

Plasmids and other reagents generated in this study will be made available to researchers by contacting. The patient-derived materials are proprietary to Prof. John S. Kuo, but available on personal requests via standard institution/university agreements.

## Code availability

The code used for differential gene expression analysis is available without restrictions at https://github.com/yuanna23/GBM_elife.

## Data availability

Source Data 1 contains all the numerical data used to generate the figures and all the original images present in the final figures.

## Supporting information

Key resources table

## Supplementary Figure and Table Legends

**Figure 1 – Figure Supplement 1. RQC pathway activity in GBM cells** (A) Immunofluorescence staining shows elevated NEMF and reduced ANKZF1 endogenous protein levels in the tumor tissue of the GBM mouse model compared to wild-type brain tissue. Tumor identification is indicated by GFP (green). (B) Quantification of A (n=3; unpaired Student’s t-test; ****, *P* < 0.0001). (C) Western blot analysis of select RQC factors in control cell lines (SVG, NHA) and GBM cell lines (SF268, GSC827), using ACTIN as the loading control. Red numbers represent fold changes in protein levels relative to controls (SVG).

**Figure 1 – Figure Supplement 2. AT repeat sequences mimicking CAT-tails induce protein aggregates in cells** (A) Western blot analysis of ATP5α in GSC and NHA cells, using GAPDH as the loading control. The purple arrowhead indicates the modified ATP5α form; “short” and “long” refer to exposure time. Red numbers represent fold changes in protein levels relative to controls (the leftmost bands); purple numbers represent fold changes in protein levels of the modified ATP5α form relative to the control (the leftmost band). (B) Western blot analysis of Flag-tagged ATP5α in GSC and control cells, using ACTIN as the loading control. The red arrowhead indicates the modified Flag-ATP5α form. (C) Immunofluorescence staining shows that Flag-tagged ATP5⍰-AT3 and ATP5⍰-AT20 (green) form aggregates in GBM and control cells, using TOM20 (red) as a mitochondrial marker. (D) Quantification of C (n=3; chi-squared test; ***, *P* < 0.001; ****, *P* < 0.0001); the total number of cells counted is indicated in the columns. (E) Western blot of Flag-tagged ATP5α-GS3 and ATP5α-GS20 in GBM cells, using ACTIN as the loading control. (F) Immunofluorescence staining shows that Flag-tagged ATP5⍰-GS3 and ATP5⍰-GS20 (green) do not form aggregates in GBM cells, using TOM20 (red) as a mitochondrial marker. (G) Quantification of F (n=3; chi-squared test; ns, not significant); the total number of cells counted is indicated in the columns.

**Figure 1 – Figure Supplement 3. Aggregation of CAT-tailed mitochondrial proteins observed *in vivo*** (A) Immunofluorescence staining shows that endogenous NDUS3 protein aggregates in GBM cells, with TOM20 (red) as a mitochondrial marker. White arrows indicate NUDS3 protein aggregates. (B) Quantification of A (n=3; chi-squared test; *, *P* < 0.05); the total number of cells counted is indicated in the columns. (C) Immunofluorescence staining reveals that endogenous ATP5⍰ protein forms aggregates in tumor tissue from the GBM mouse model, but not in wild-type brain tissue, using TOM20 (blue) as a mitochondrial marker. Tumor identification is indicated by GFP (green). White arrowheads indicate ATP5⍰ (red) aggregates. Yellow lines indicate the regions selected for intensity analysis in (D). (D) Fluorescence intensity profiles show the signals of ATP5⍰ (red) and TOM20 (blue) in wild-type and tumor tissues. Black arrows indicate ATP5⍰ aggregates located outside of mitochondria. (E) Quantification of C (n=3; chi-squared test; ****, *P* < 0.0001); the total number of cells counted is indicated in the columns.

**Figure 2 – Figure Supplement 1. Aberrant mitochondrial function in GBM cells** (A) JC-10 staining reveals elevated mitochondrial membrane potentials in GBM cells compared to NHA (control) cells (n=3; unpaired Student’s t-test; ***, *P* < 0.001). (B) Analysis with BioTracker ATP red dye staining shows reduced mitochondrial ATP production in GBM cells compared to NHA (control) cells, using MitoTracker Green as the mitochondrial mass indicator for normalization. (C) Quantification of (B) (n=3; unpaired Student’s t-test; ****, *P* < 0.0001). (D) Western blot of NEMF and ANKZF1 in GBM and control cells, confirming the successful overexpression and knockdown of target proteins, using ACTIN as the loading control. (E) JC-10 staining reveals no change of mitochondrial membrane potential in GBM cells, upon overexpression of ATP5⍰-GS3 and ATP5⍰-GS20 (n=3; unpaired Student’s t-test; ****, *P* < 0.0001; ns, not significant).

**Figure 3 – Figure Supplement 1. Cycloheximide does not impact mitochondrial functions** (A) MPTP activity assay shows that MPTP opening is not affected by the cycloheximide treatment (100 µg/mL) in cells. (B) Quantification of (A) (n=3; unpaired Student’s t-test; ns, not significant). (C) Immunofluorescence staining reveals no inhibition of endogenous ATP5⍰ protein aggregation by cycloheximide (100 µg/mL) treatment in GBM cells, using TOM20 (red) as a mitochondrial marker. (D) Quantification of (C) (n=3; chi-squared test; ns, not significant); the total number of cells counted is indicated in the columns. (E) MPTP activity assay reveals the increased Calcien signal in GBM cells, upon overexpression of ATP5⍰-AT3 and ATP5⍰-AT20 with concurrent genetic inhibition of the msiCAT-tailing pathway (n=3; unpaired Student’s t-test; **, *P* < 0.01; ***, *P* < 0.001; ****, *P* < 0.0001).

**Figure 3 – Figure Supplement 2. The CAT-tailed ATP5IZ variant has no interaction with MPTP proteins** (A) Calcium retention capacity (CRC) assay of isolated mitochondria, measured with Calcium Green-5N dye, upon cycloheximide (100 µg/mL) treatment and CAT-tailing enhancement (oeNEMF and siANKZF1). (B) Statistic of (A) shows changes in CRC in GBM cells or control cells (n=10; unpaired Student’s t-test; **, *P* < 0.01; ***, *P* < 0.001). (C) Co-immunoprecipitation data show no direct interaction between ATP5⍰ and either CypD or ANT1/2 can be found in GBM cells. Red arrowheads indicate target proteins. (D) Western blotting of cytosolic and isolated mitochondrial fractions shows ATP5⍰-AT3 expression reduces CypD levels in GBM cells, using TOM20 as a mitochondrial marker and loading control.

**Figure 4 – Figure Supplement 1. Effect of GS repeat tails on GBM proliferation and migration** (A) MTT assay indicates no significant change in GBM proliferation upon ATP5⍰-GS3 and ATP5⍰-GS20 expression (n=3; unpaired Student’s t-test; **, *P* < 0. 01; ns, not significant). (B) Wound-healing assay reveals enhanced GBM migration upon ATP5⍰-AT3 and ATP5⍰-AT20 expression. (C) Quantification of (B) shows an increased healing rate, indicated by scratch wound coverage at both 24 and 48 hours (n=3; unpaired Student’s t-test; **, *P* < 0.01). (D) Transwell assay reveals no significant alteration in GBM migration upon ATP5⍰-GS3 and ATP5⍰-GS20 expression. (E) Quantification of (D) shows the number of migrated cells (n=3; unpaired Student’s t-test; ns, not significant). (F) qRT-PCR reveals no increase in mRNA levels of mitochondrial unfolded protein response genes, as normalized to *ACTB* as the control (n=4; unpaired Student’s t-test; **, *P* < 0.01; ns, not significant).

**Figure 4 – Figure Supplement 2.** GBM cells exhibit increased resistance to apoptosis (A) TUNEL staining shows that GBM cells are more resistant to staurosporine (STS, 1 µM)-induced apoptosis compared to control cells, using TUNEL-Cy3 as an apoptotic cell indicator and DAPI as a nucleus indicator. (B) Quantification of A shows the percentage of TUNEL-positive cells in the population (n=3; unpaired Student’s t-test; ***, *P* < 0.0001; ****, *P* < 0.0001), using DMSO as the vehicle control. (C) Western blot analysis of PARP shows that GBM cells are more resilient against STS-induced apoptosis at 30-, 90-, and 180-min post-treatment. Cleaved PARP is used as an apoptosis marker. ACTIN and GAPDH are used as loading controls. Red numbers below each blot represent the ratios of cleaved PARP (c-PARP) to total PARP protein. (D, F) Flow cytometry analysis using Annexin V-FITC/Propidium Iodide (PI) staining shows alterations in apoptosis rates in GBM cells upon ATP5⍰-AT3, ATP5⍰-AT20, ATP5⍰-GS3, and ATP5⍰-GS20 expression. The apoptotic cell population (Annexin V positive, PI negative) is represented in the fourth quadrant (right lower). (E, G) Quantification of (D, F) shows the percentages of apoptotic cells (n=3; unpaired Student’s t-test; **, *P* < 0.001; ***, *P* < 0.0001).

**Figure 5 – Figure Supplement 1. No effect of Cycloheximide on GBM apoptosis response** (A) Caspase-3/7 activity assay shows increased apoptosis in GBM cells caused by anisomycin treatment (n=3; unpaired Student’s t-test; ***, *P* < 0.001; ****, *P* <0.0001; compared to the control group (DMSO) at the corresponding time). (B, C) Western blot analysis of PARP in anisomycin-treated and cycloheximide-treated GSC cells indicates that pharmacological inhibition of the msiCAT-tailing pathway enhances STS-induced apoptosis, using ACTIN as a loading control. Red numbers below each blot represent the ratios of cleaved PARP (c-PARP) to total PARP protein. (D, F) Flow cytometry analysis using Annexin V-FITC/Propidium Iodide (PI) staining shows alterations in apoptosis rates in GBM cells upon genetic (D) and pharmacological (F) inhibition of the msiCAT-tailing pathway. The apoptotic cell population (Annexin V positive, PI negative) is represented in the fourth quadrant (right lower). (E, G) Quantification of (D, F) shows the percentages of apoptotic cells (n=3; unpaired Student’s t-test; **, *P* < 0.01; ***, *P* < 0.001; ****, *P* <0.0001; ns, not significant).

## Acknowledgements

The patient-derived GSC and NSC lines were a gift from Dr. John S. Kuo at the University of Texas, Austin. We are also grateful to Dr. Chunzhang Yang at the National Cancer Institute and Dr. Russell O. Pieper at the University of California, San Francisco, for the cell lines. We thank Dr. Hoshino at Nagoya City University and Dr. Inada at Tohoku University for providing the plasmids. We also thank the members of the Wu lab at Southern Methodist University and the members of the Lu lab at UCSF for their assistance and discussions.

