## Supplementary figures and images for "Mitochondrial Protein Carboxyl-Terminal Alanine-Threonine Tailing Promotes Human Glioblastoma Growth by Regulating Mitochondrial Function"

### Supplementary Figures

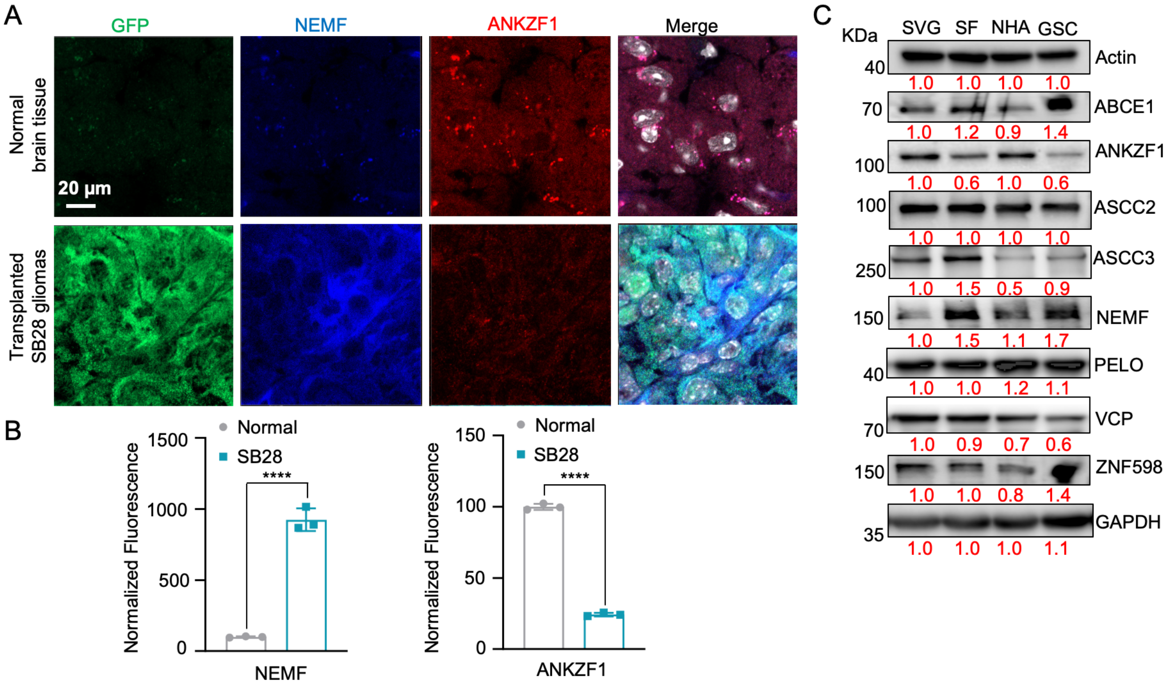

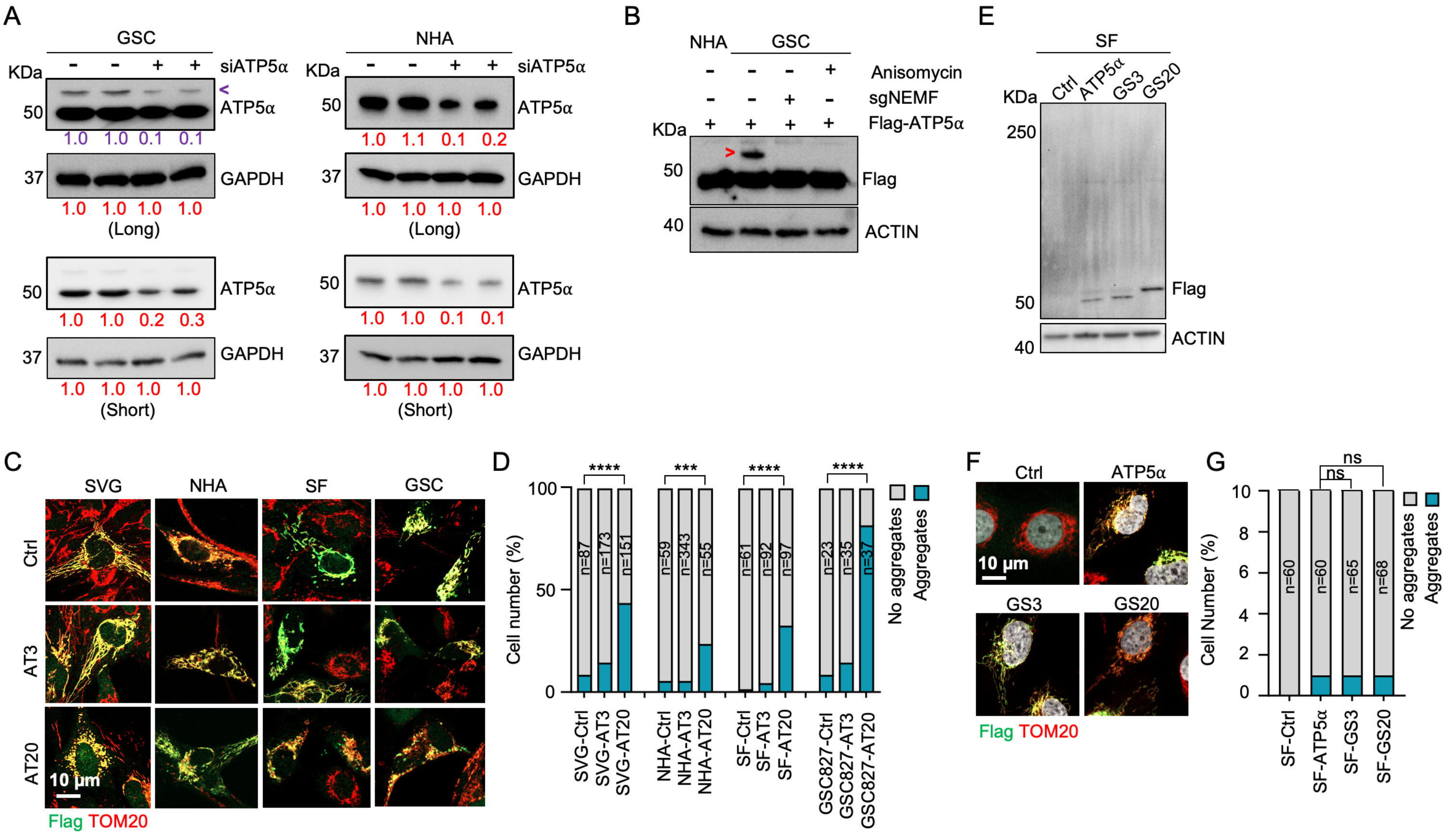

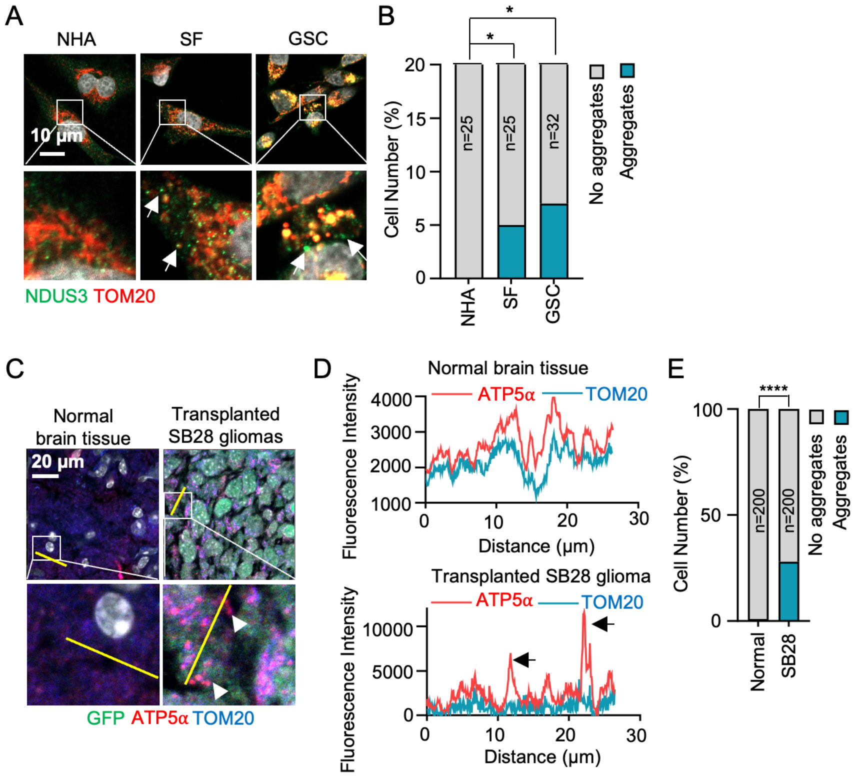

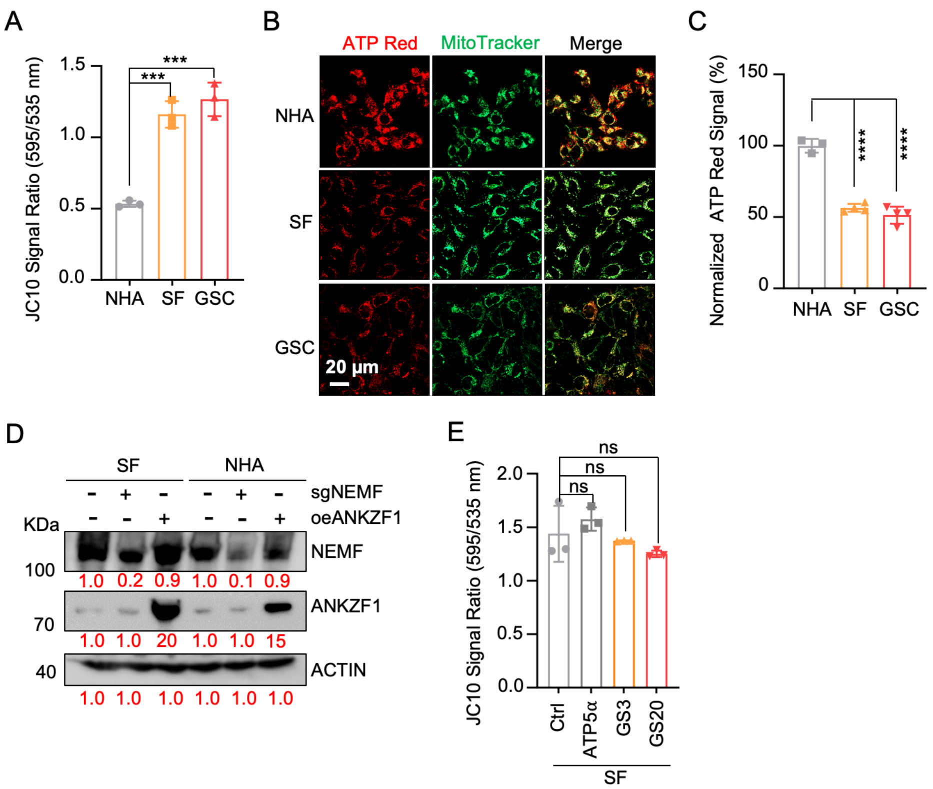

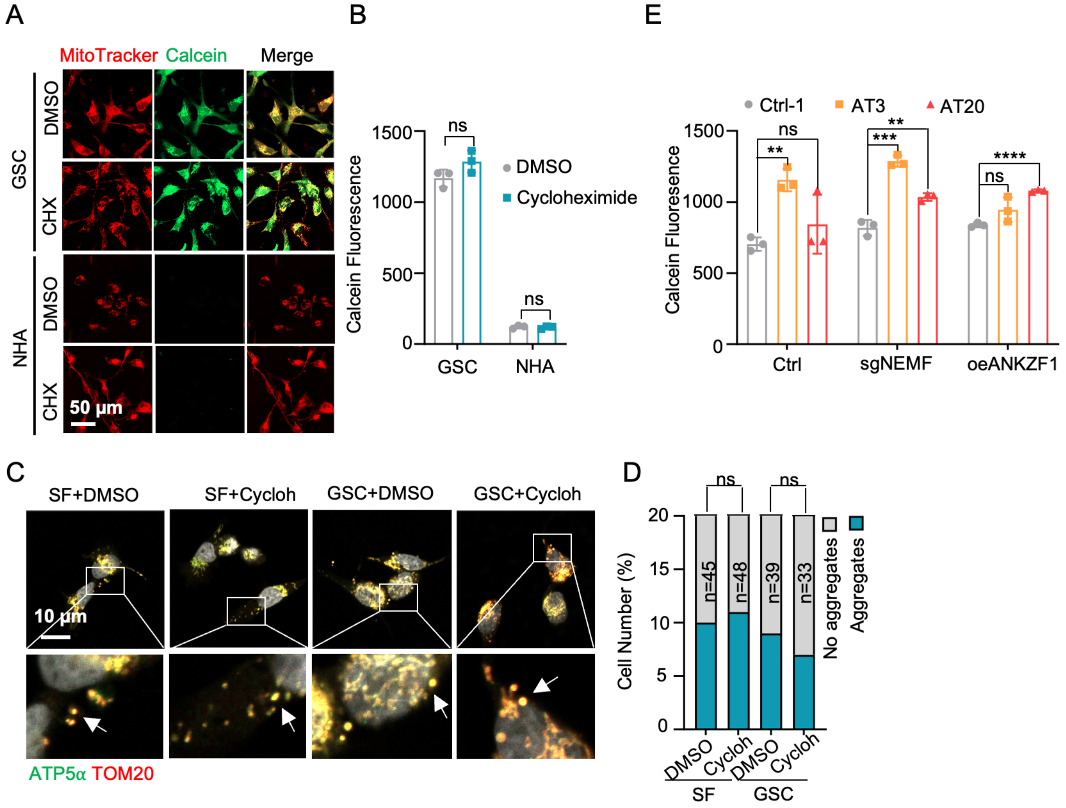

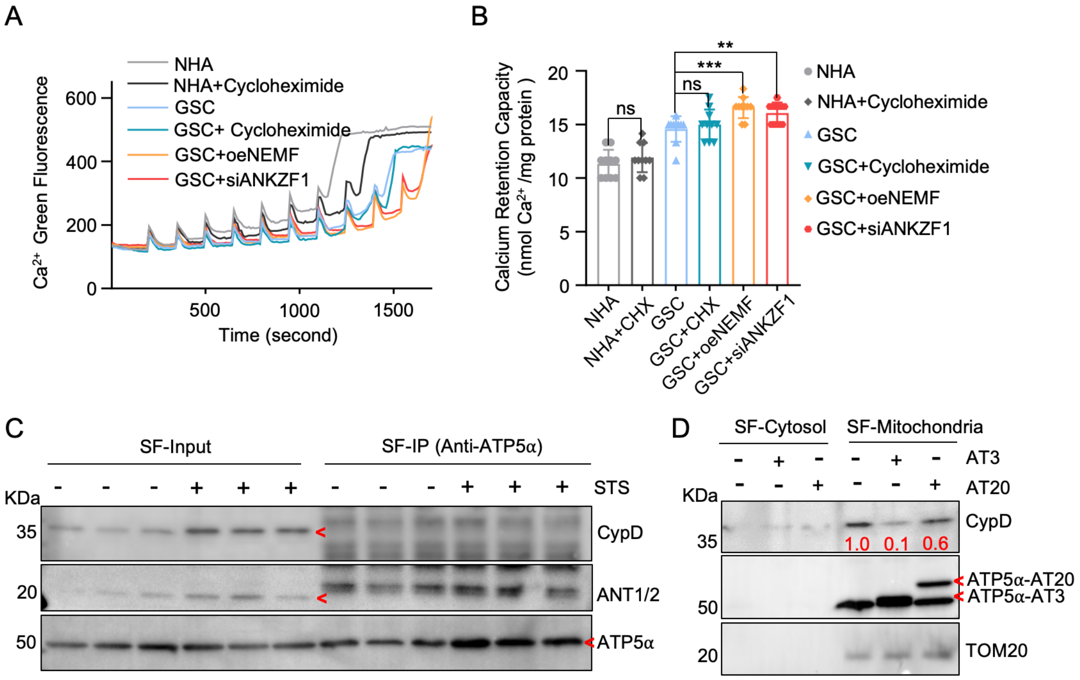

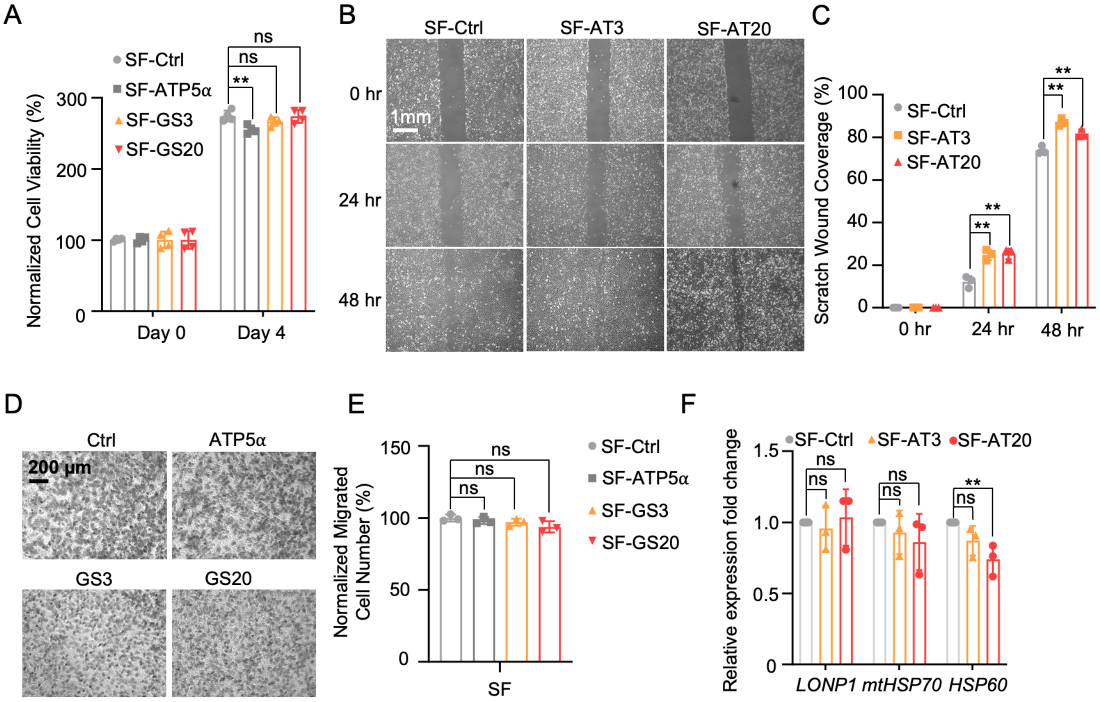

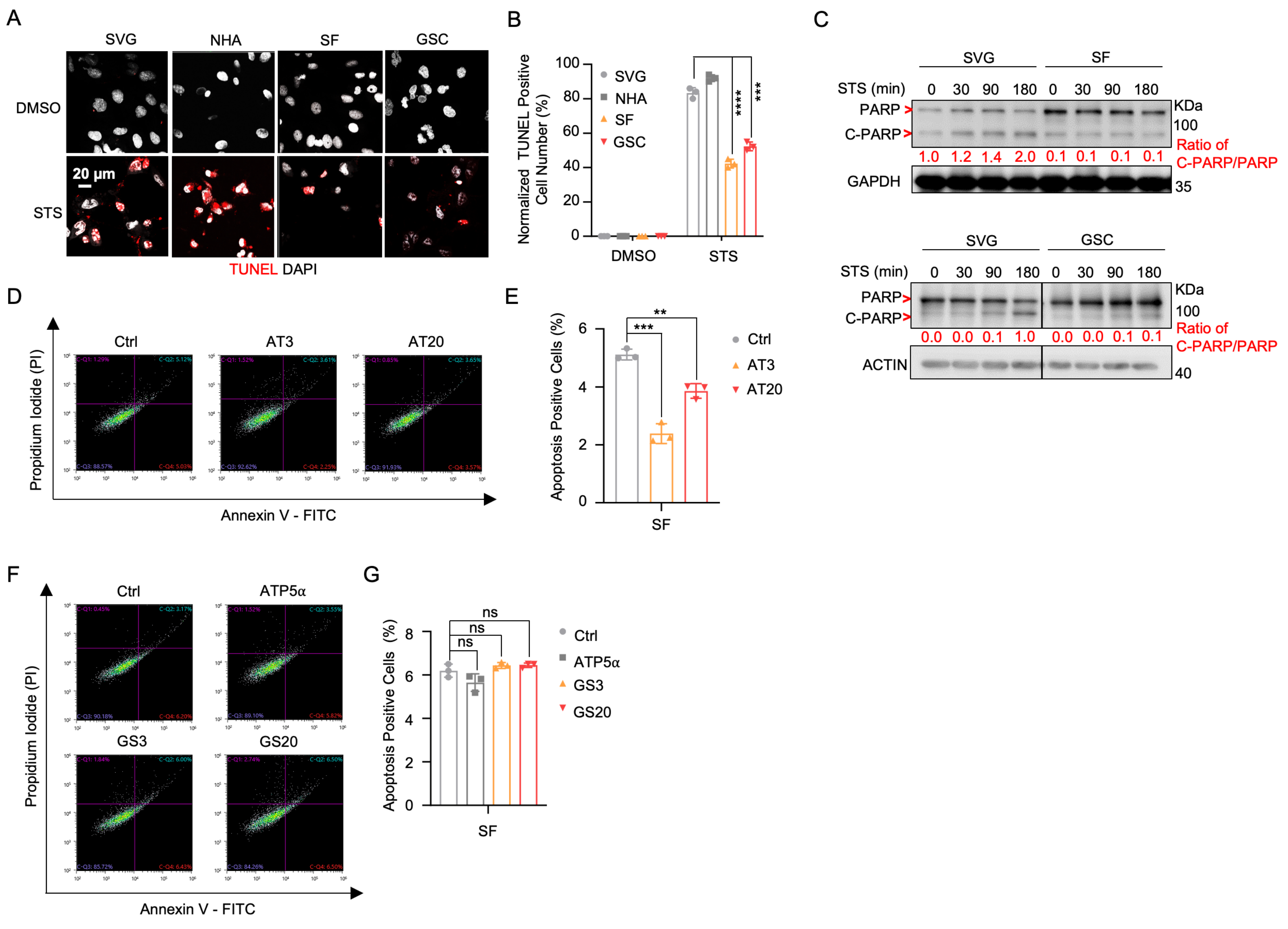

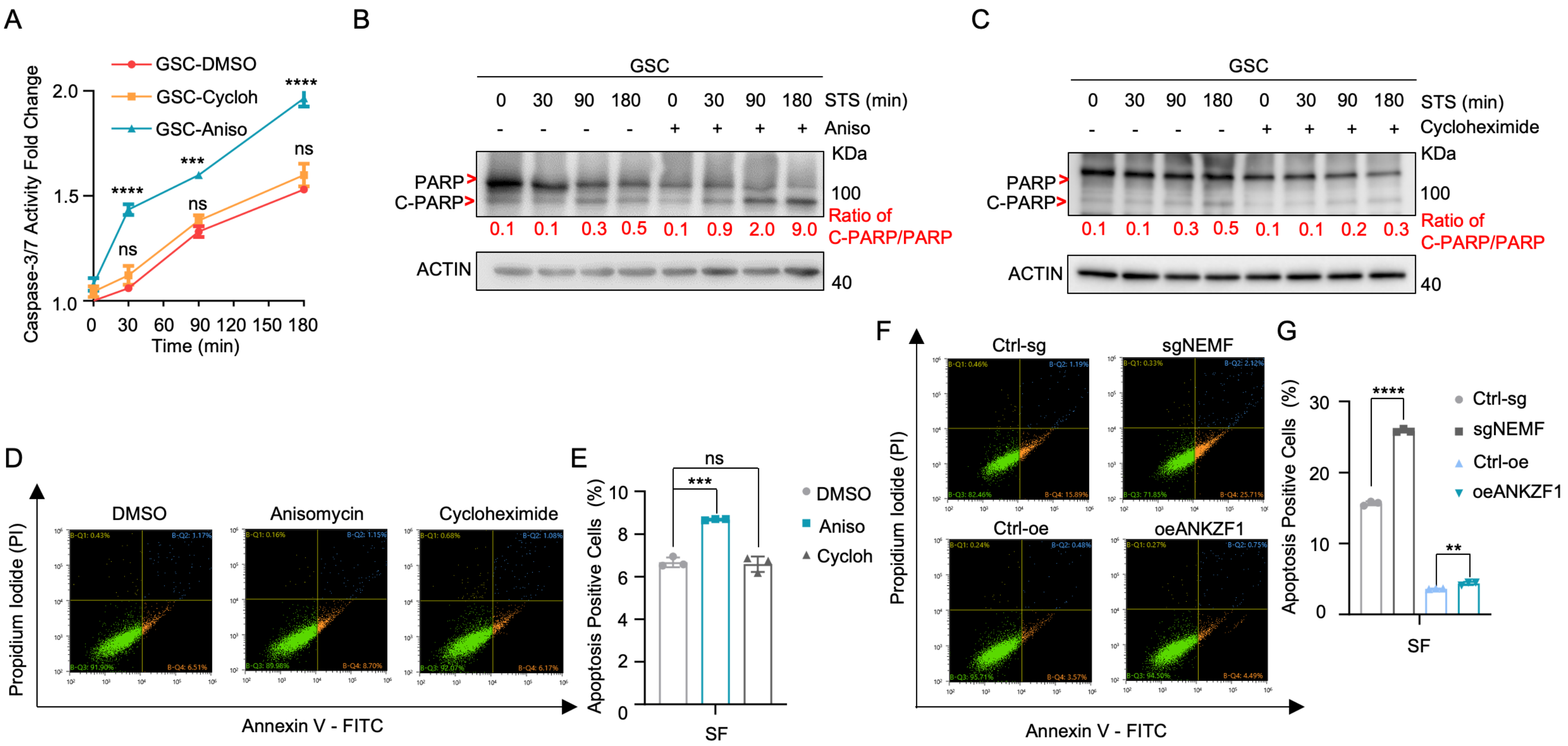
