## Supplementary material for "Mitochondrial Protein Carboxyl-Terminal Alanine-Threonine Tailing Promotes Human Glioblastoma Growth by Regulating Mitochondrial Function": Key resources table

| **Key Resources Table** | | | | |
| --- | --- | --- | --- | --- |
| **Reagent type (species) or resource** | **Designation** | **Source or reference** | **Identifiers** | **Additional information** |
| strain, strain background (mice) | C57BL/6J mice | Dr. Rongze Olivia Lu | RRID: IMSR_JAX:000664 |  |
| cell line (*Homo-sapiens*) | SVG p12 | ATCC | RRID:  CRL-8621 |  |
| cell line (*Homo-sapiens*) | SF268 | Dr. Rongze Olivia Lu | RRID: CVCL_1689 |  |
| cell line (*Homo-sapiens*) | GL261 | Dr. Rongze Olivia Lu | RRID: CVCL_Y003 |  |
| cell line (*Homo-sapiens*) | SB28 | Dr. Rongze Olivia Lu | RRID: CVCL_A5ED |  |
| cell line (*Homo-sapiens*) | GSC827 | Dr. Chun-Zhang Yang | N/A | Patient-derived |
| cell line (*Homo-sapiens*) | NSC | Dr. John S Kuo | N/A | Human-derived |
| cell line (*Homo-sapiens*) | NSC26 | Dr. John S Kuo | N/A | Human-derived |
| cell line (*Homo-sapiens*) | GSC33 | Dr. John S Kuo | N/A | Patient-derived |
| cell line (*Homo-sapiens*) | GSC22 | Dr. John S Kuo | N/A | Patient-derived |
| cell line (*Homo-sapiens*) | GSC99 | Dr. John S Kuo | N/A | Patient-derived |
| cell line (*Homo-sapiens*) | GSC105 | Dr. John S Kuo | N/A | Patient-derived |
| cell line (*Homo-sapiens*) | GSC107 | Dr. John S Kuo | N/A | Patient-derived |
| cell line (*Homo-sapiens*) | NHA | Dr. Russell O. Pieper | RRID: CVCL_E3G5 |  |
| transfected construct (human) | pLV[CRISPR]-hCas9:T2A:Neo-U6>Scramble [gRNA#1] | This study;  VectorBuilder | Cat#: VB240227-1635qjy | Lentiviral construct to express sgRNA |
| transfected construct (human) | pLV[CRISPR]-hCas9:T2A:Neo-U6>hNEMF [gRNA#1579] | This study;  VectorBuilder | Cat#: VB900124-2190daq | Lentiviral construct to express sgRNA |
| transfected construct (human) | pLV[Exp]-Bsd-EF1A>ORF_Stuffer | This study;  VectorBuilder | Cat#: VB900145-3633yjp | Lentiviral construct to express target gene |
| transfected construct (human) | pLV[Exp]-EGFP:T2A:Puro-EF1A>mCherry | This study;  VectorBuilder | Cat#: VB010000-9298rtf | Lentiviral construct to express target gene |
| transfected construct (human) | pLV[Exp]-Bsd-EF1A>hANKZF1 [NM_001042410.2]/HA | This study;  VectorBuilder | Cat#: VB240227-1626epe | Lentiviral construct to express target gene |
| transfected construct (human) | pLV[Exp]-mCherry/Neo-EF1A>hANKZF1 [NM_001042410.2] | This study;  VectorBuilder | Cat#: VB900124-2193gcv | Lentiviral construct to express target gene |
| antibody | anti-COX4 (Rabbit polyclonal) | Abcam | Cat#: ab209727,  RRID: AB_3717302 | WB (1:1000) |
| antibody | anti-β-Actin [C4] (Mouse monoclonal) | Santa Cruz | Cat#: sc-47778,  RRID: AB_626632 | WB (1:1000) |
| antibody | anti-Flag (Mouse monoclonal) | Millipore Sigma | Cat#: F1804,  RRID: AB_262044 | WB (1:1000) |
| antibody | anti-ANT1/2 (Rabbit polyclonal) | Proteintech | Cat#: 17796-1-AP,  RRID: AB_2190358 | WB (1:1000) |
| antibody | anti-CypD (Rabbit polyclonal) | Proteinetch | Cat#: 15997-1-AP,  RRID: AB_2190199 | WB (1:1000) |
| antibody | anti-ATP5a (Rabbit polyclonal) | Cell Signaling Technology | Cat#: 18023,  RRID: AB_2687556 | WB (1:1000)  IF (1:500) |
| antibody | anti-PARP1 (Rabbit polyclonal) | Abclonal | Cat#: A0942,  RRID: AB_2757470 | WB (1:1000) |
| antibody | anti-GAPDH (Rabbit polyclonal) | Abclonal | Cat#: A19056,  RRID: AB_2862549 | WB (1:1000) |
| antibody | anti-TOMM20 (Mouse monoclonal) | Santa Cruz | Cat#: sc-17764,  RRID: AB_628381 | WB (1:1000)  IF (1:500) |
| antibody | anti-MTCO2 (Rabbit polyclonal) | Proteintech | Cat#: 55070-1-AP,  RRID: AB_10859832 | WB (1:1000)  IF (1:500) |
| antibody | anti-NDUS3 (Mouse monoclonal) | Abcam | Cat#: ab14711,  RRID: AB_301429 | WB (1:1000)  IF (1:1000) |
| antibody | anti-NEMF (Rabbit polyclonal) | Proteintech | Cat#: 11840-1-AP,  RRID: AB_2183413 | WB (1:1000)  IF (1:500) |
| antibody | anti-ANKZF1 (Mouse monoclonal) | Santa Cruz | Cat#: sc-398713,  RRID: AB_3094545 | WB (1:1000)  IF (1:500) |
| antibody | anti-ATP5a (Mouse monoclonal) | Abcam | Cat#: ab14748,  RRID: AB_301447 | WB (1:1000)  IF (1:500) |
| antibody | anti-TOMM20 (Rat monoclonal) | Abcam | Cat#: Ab289670,  RRID: AB_3097753 | WB (1:1000)  IF (1:500) |
| antibody | anti-GFP (Chicken polyclonal) | Abcam | Cat#: Ab13970,  RRID: AB_300798 | WB (1:1000)  IF (1:500) |
| antibody | anti-rabbit HRP (Goat polyclonal) | Invitrogen | Cat#: G21234,  RRID: AB_2536530 | WB (1:5000) |
| antibody | anti-mouse HRP (Goat polyclonal) | Invitrogen | Cat#: PI31430,  RRID: AB_228307 | WB (1:5000) |
| antibody | anti-Mouse IgG (H+L) Highly Cross-Adsorbed Secondary Antibody, Alexa Fluor 488 (Goat polyclonal) | Invitrogen | Cat#: A32723,  RRID: AB_2633275 | IF (1:300) |
| antibody | anti-Rabbit IgG (H+L) Cross-Adsorbed Secondary Antibody, Alexa Fluor 633 (Goat polyclonal) | Invitrogen | Cat#: A21071,  RRID: AB_2535732 | IF (1:300) |
| antibody | anti-Rabbit IgG (H+L) Highly Cross-Adsorbed Secondary Antibody, Alexa Fluor 568 (Goat polyclonal) | Invitrogen | Cat#: A11036,  RRID: AB_10563566 | IF (1:300) |
| recombinant DNA reagent | pcDNA3.1+/C-(K)-DYK-ATP5F1A | This study;  GenScript | Clone ID:  OHu25769D |  |
| recombinant DNA reagent | pcDNA3.1+/C-(K)-DYK-ATP5F1A-AT3 | This study;  GenScript | N/A |  |
| recombinant DNA reagent | pcDNA3.1+/C-(K)-DYK-ATP5F1A-AT20 | This study;  GenScript | N/A |  |
| recombinant DNA reagent | pcDNA3.1+/C-(K)-DYK-ATP5F1A-GS3 | This study;  GenScript | N/A |  |
| recombinant DNA reagent | pcDNA3.1+/C-(K)-ATP5F1A-DYK-GS20 | This study;  GenScript | N/A |  |
| recombinant DNA reagent | pCMV-5×FLAG-β-globin-control | Dr. Hoshino and Dr. Inada | N/A |  |
| recombinant DNA reagent | pCMV-5×FLAG-β-globin-non-stop | Dr. Hoshino and Dr. Inada | N/A |  |
| recombinant DNA reagent | pCMV6-DDK-NEMF (NM_004713) | ORIGENE | Cat#: RC216806L3 |  |
| sequenced-based reagent | lonp1_F | This study;  GeneWiz | PCR primers | TGCCTTGAACCCTCTCTAC |
| sequenced-based reagent | lonp1_R | This study;  GeneWiz | PCR primers | TCTGCTTGATCTTCTCCTCC |
| sequenced-based reagent | mthsp70_F | This study;  GeneWiz | PCR primers | ACTCCTCCATTTATCCGCC |
| sequenced-based reagent | mthsp70_R | This study;  GeneWiz | PCR primers | ACCTTTGCTTGTTTACCTTCC |
| sequenced-based reagent | hsp60_F | This study;  GeneWiz | PCR primers | ACCTGCTCTTGAAATTGCC |
| sequenced-based reagent | hsp60_R | This study;  GeneWiz | PCR primers | CAATCCCTCTTCTCCAAACAC |
| sequenced-based reagent | actb_F | This study;  GeneWiz | PCR primers | TGTTTGAGACCTTCAACACC |
| sequenced-based reagent | actb_R | This study;  GeneWiz | PCR primers | ATGTCACGCACGATTTCC |
| commercial assay or kit | AGM^TM^ SingleQuots^TM^ Supplements | Lonza | Cat#: CC-4123 |  |
| commercial assay or kit | MTT assay kit | Roche | Cat#: 11465007001 |  |
| commercial assay or kit | Seahorse Cell Mito Stress Test kit | Agilent | Cat#: 103010-100 |  |
| commercial assay or kit | NativePAGE Running Buffer Kit | Invitrogen | Cat#: BN2007 |  |
| commercial assay or kit | NativePAGE Sample Prep Kit | Invitrogen | Cat#: BN2008 |  |
| commercial assay or kit | TUNEL assay | ApexBio | Cat#: K1134 |  |
| commercial assay or kit | Annexin V-FITC/PI apoptosis assay | BioLegend | Cat#: 640914 |  |
| commercial assay or kit | Seahorse XF DMEM medium | Agilent | Cat#: 103575-100 |  |
| commercial assay or kit | Mitochondrial Transition Pore Assay | Invitrogen | Cat#: I35103 |  |
| chemical compound, drug | DMEM | ATCC | Cat#: 302002 |  |
| chemical compound, drug | FBS | Biowest | Cat#: S1620-100 |  |
| chemical compound, drug | Penicillin/streptomycin | Gibco | Cat#: 15140122 |  |
| chemical compound, drug | G418 | Gibco | Cat#: 10131027 |  |
| chemical compound, drug | 0.25% trypsin solution | ATCC | Cat#: SM2003C |  |
| chemical compound, drug | ABM^TM^ Basal Medium | Lonza | Cat#: CC-3187 |  |
| chemical compound, drug | Accutase | Corning | Cat#: 25058CI |  |
| chemical compound, drug | Neural basal-A Medium | Gibco | Cat#: 10888022 |  |
| chemical compound, drug | B27 | Gibco | Cat#: 17504044 |  |
| chemical compound, drug | N2 | Gibco | Cat#: 17502048 |  |
| chemical compound, drug | EGF and FGF | Shenandoah Biotech | Cat#: PB-500-017 |  |
| chemical compound, drug | Antibiotic-Antimycotic | Gibco | Cat#: 15240062 |  |
| chemical compound, drug | L-Glutamine | Gibco | Cat#: 250300810 |  |
| chemical compound, drug | Geltrex | Thermo Fisher | Cat#: A1413202 |  |
| chemical compound, drug | X-tremeGENE | Sigma | Cat#: 6366244001 |  |
| chemical compound, drug | Anisomycin | Fisher Scientific | Cat#: AAJ62964MF |  |
| chemical compound, drug | Cycloheximide | Fisher Scientific | Cat#: AC357420010 |  |
| chemical compound, drug | Temozolomide | Millipore Sigma | Cat#:  50-060-90001 |  |
| chemical compound, drug | Formaldehyde | Thermo Fisher | Cat#: BP531-500 |  |
| chemical compound, drug | Triton X-100 | Thermo fisher | Cat#: T9284 |  |
| chemical compound, drug | Lipofectamine 3000 | Invitrogen | Cat#: L3000015 |  |
| chemical compound, drug | Normal goat serum | Jackson Immuno | Cat#:  005-000-121 |  |
| chemical compound, drug | DAPI | Thermo Fisher | Cat#: 57-481-0 |  |
| chemical compound, drug | Fluoromount-G Anti-Fade | Southern Biotech | Cat#: 0100-35 |  |
| chemical compound, drug | Puromycin | ARCOS organics | Cat#: 227420100 |  |
| chemical compound, drug | Protease inhibitor | Bimake | Cat#: B14002 |  |
| chemical compound, drug | Bradford | BioVision | Cat#:  K813-5000-1 |  |
| chemical compound, drug | Mannitol | Fisher Scientific | Cat#: AA3334236 |  |
| chemical compound, drug | Sucrose | Fisher Scientific | Cat#: AA36508A1 |  |
| chemical compound, drug | HEPES | Fisher Scientific | Cat#: 15630106 |  |
| chemical compound, drug | Western Lightning Plus-ECL | PerkinElmer Inc. | Cat#: NEL104001EA |  |
| chemical compound, drug | 4-12% Tris-Glycine gel | Invitrogen | Cat#: WXP41220BOX |  |
| chemical compound, drug | PVDF membrane | Millipore | Cat#: ISEQ00010 |  |
| chemical compound, drug | EGTA | Fisher Scientific | Cat#: 28-071-G |  |
| chemical compound, drug | Digitonin | Thermo Fisher | Cat#: BN2006 |  |
| chemical compound, drug | G-250 | GoldBio | Cat#: C-460-5 |  |
| chemical compound, drug | 3-12% Bis-Tris Native gel | Invitrogen | Cat#: BN1001BOX |  |
| chemical compound, drug | Acetic acid | Thermo Fisher | Cat#: 9526-33 |  |
| chemical compound, drug | TMRM | Invitrogen | Cat#: I34361 |  |
| chemical compound, drug | JC-10 | AdipoGen | Cat#:  50-114-6552 |  |
| chemical compound, drug | Succinate | Thermo Fisher | Cat#: 041983.A7 |  |
| chemical compound, drug | Hank's Balanced Salt Solution | Thermo Fisher | Cat#: 14025092 |  |
| chemical compound, drug | Calcium green-5N | Invitrogen | Cat#: C3737 |  |
| chemical compound, drug | Cyclosporine A | Thermo Fisher | Cat#: AC457970010 |  |
| chemical compound, drug | Ethanol | Thermo Fisher | Cat#: R40135 |  |
| chemical compound, drug | Crystal violet | Sigma | Cat#: V5265 |  |
| chemical compound, drug | Proteinase K | Invitrogen | Cat#: 25530049 |  |
| chemical compound, drug | Caspase‑3/7 detection reagents | Invitrogen | Cat#: C10432 |  |
| chemical compound, drug | ATP‐red dye | Millipore | Cat#: SCT045 |  |
| chemical compound, drug | MitoTracker-Green | Invitrogen | Cat#: M7514 |  |
| chemical compound, drug | Protein A/G magnetic beads | Pierce | Cat#: 88802 |  |
| chemical compound, drug | M.O.M. blocking reagent | Vector Laboratories | Cat#: BMK-2202 |  |
| software, algorithm | SPSS | SPSS | RRID: [SCR_002865](https://scicrunch.org/resolver/SCR_002865) |  |
| software, algorithm | BioRender | BioRender | RRID: SCR_018361 | https://biorender.com/ |
| software, algorithm | GraphPad Prism 9.4.1 | GraphPad | RRID: SCR_002798 | https://www.graphpad.com/scientific-software/prism/ |
| software, algorithm | ImageJ 1.53t | NIH | RRID: SCR_003070 | https://imagej.nih.gov/ij/download.html |
| software, algorithm | ZEN (blue edition) | ZEISS | RRID: SCR_013672 | https://www.zeiss.com/microscopy/us/products/microscope-software.html |
| software, algorithm | Gen5 | Agilent Technologies (BioTek) | RRID: SCR_017317 | https://www.biotek.com/products/software-robotics-software/gen5-microplate-reader-and-imager-software/ |
| software, algorithm | Endnote 20 | Clarivate | RRID: SCR_014001 | https://endnote.com/downloads |
